# Insights into early cochlear damage induced by potassium channel deficiency

**DOI:** 10.1101/2025.04.08.647809

**Authors:** E. Rías, C. Carignano, V. C. Castagna, L. Dionisio, J. A. Ballestero, G. Paolillo, I. Ouwerkerk, M. E. Gomez-Casati, G. Spitzmaul

## Abstract

Hearing loss (HL) is the most common sensory disorder, caused by genetic mutations and acquired factors like presbycusis and noise exposure. A critical factor in HL development is the dysfunction of potassium (K^+^) channels, essential for sensory cell function in the organ of Corti (OC). Inner and outer hair cells (IHCs and OHCs) convert sound into electrical signals, while supporting cells (SCs) maintain ionic and structural balance. KCNQ4 channels, located in the basal membrane of OHCs, regulate K^+^ efflux. Mutations in KCNQ4 are linked to progressive HL (DFNA2), noise-induced hearing loss, and presbycusis, leading to K^+^ accumulation, cellular stress, and OHC death. Gene editing or pharmacological activation of KCNQ4 has shown potential in partially preventing HL in mouse models. In this study, we demonstrate KCNQ4 deletion disrupts the localization of key proteins like prestin and BK channels, alters OHC organization, and induces apoptosis in sensory and SC. Spiral ganglion neurons (SGNs) also degenerate over time. Despite these structural changes, noise exposure does not exacerbate OHC damage in our KCNQ4-deficient model. This highlights KCNQ4’s role in maintaining ion homeostasis and cochlear function, as its absence triggers widespread dysfunction in the OC. The present study demonstrates that disruptions in a single cell type can have a cascade effect on overall cochlear health. Understanding the molecular and cellular consequences of KCNQ4 mutations is crucial for developing targeted therapies to mitigate progressive HL caused by genetic and environmental factors.

**Highlights:**

- Hearing function is altered in KCNQ4 KO animals from young ages.
- Inner hair cells show structural alterations before their death.
- KCNQ4 absence impairs membrane localization of key functional proteins.
- Hair cell and neuron loss is mediated by apoptosis.
- Supporting cells and satellite cells also contribute to tissue degeneration in KCNQ4 KO animals
- Noise exposure does not exacerbate hair cell damage in KCNQ4-KO mice.

## 1. INTRODUCTION

Hearing loss (HL) is the most prevalent sensory disorder, affecting millions globally and significantly diminishing quality of life [1]. The primary causes of hearing impairment include both genetic mutations and acquired factors, such as presbycusis (age-related hearing loss) and noise-induced hearing loss (NIHL) [2]. These two types of HL are frequently associated with the dysfunction or loss of potassium (K^+^) channels, which are vital for K^+^ recycling in the organ of Corti (OC) [3]. The OC is composed of sensory cells known as inner hair cells (IHCs) and outer hair cells (OHCs), which are responsible for converting sound vibrations into electrical signals. Supporting cells (SCs) provide structural scaffolding for the OC, protecting the hair cells (HCs) while also regulating the ionic composition of the perilymph to maintain proper HC function [4]. The connection between the OC and the central nervous system is mediated by the spiral ganglion neurons (SGNs), bipolar neurons with peripheral axons that innervate sensory hair cells in the cochlea and central axons to project to the cochlear nucleus. SGN fibers transmit the receptor potentials generated by IHCs to their central projections, forming the auditory nerve that conveys signals to the brain. The generation of these receptor potentials are mostly dependent on K^+^ influx from the endolymph, as K^+^ is the primary ion responsible for HC depolarization [5]. For this reason, K^+^ channels and transporters are essential for normal hearing function.

Among the K^+^ channels expressed in the OC, KCNQ4 plays a critical role in maintaining the membrane potential of OHCs, necessary for their role in sound amplification. This channel is predominantly expressed in the basal membrane of the OHCs, where it facilitates K^+^ efflux [6–8]. Mutations that impair channel function or reduce the expression of KCNQ4 lead to non-syndromic progressive HL DFNA2, typically affecting high-frequency hearing first and worsening over time [5, 9, 10]. Such alterations result in K^+^ accumulation and chronic depolarization of OHCs, leading to cellular stress and OHC death. Genetic variants of KCNQ4 have been associated not only to DFNA2 but also to NIHL and presbycusis [3, 11–13]. In fact, *in vivo* gene editing in mice carrying pathological *Kcnq4* gene mutations, or pharmacological treatment with a KCNQ4 channel activator in a mouse model of presbycusis has been shown to partially prevent HL [14–16]. In KCNQ4 knock-out (KO) mice, OHCs are lost early on, and as the mice age, IHCs and spiral ganglion neurons (SGN) also degenerate, suggesting that the absence of KCNQ4 leads to long-term structural changes in the cochlea [6, 10].

OHC function relies not only on the presence of specific proteins but also on their precise localization within the membrane [7, 17–19]. Prestin, located in the lateral membrane, is responsible for driving electromotility, while KCNQ4, BK channels and the nicotinic acetylcholine receptor α9/α10 are localized in the basal membrane, where they play key roles in regulating ion homeostasis and efferent modulation, respectively [20]. Disruptions in this membrane compartmentalization impair both OHC function and tissue organization, as has been observed with prestin mutations [17, 21–23].

SCs surround the sensory HCs and are specialized for maintaining both structural integrity and ion balance [4, 24]. Deiters’ cells (DCs), in particular, engulf OHCs and play a vital role in maintaining extracellular K^+^ and glutamate homeostasis [4, 25]. K^+^ extruded by OHCs through KCNQ4 into the perilymph must be absorbed by DCs and transported back to the stria vascularis [24]. To achieve this, SCs express proteins involved in K^+^ ion uptake, such as Kir4.1, KCC4 and gap junction channels, such as connexins [5, 26–28]. The interconnectivity of cells within the OC means that alterations in the expression or function of critical proteins in one cell type can disrupt the function of neighboring cells, triggering a cascade of dysfunction throughout the cochlear system [10, 29–32].

A key factor in the development of HL is cellular stress, such as chronic depolarization, which contributes to the degeneration of cochlear sensory cells and neurons [10]. Cellular stress in the cochlea is often triggered by a variety of harmful conditions [33, 34]. Apoptosis plays a significant role in cochlear cell loss across various forms of HL. Although SCs exhibit greater resistance to certain types of damage compared to HCs, apoptosis has been observed in both cell types under conditions such as acoustic trauma [35, 36], ageing [37] exposure to ototoxic drugs [38], sepsis-related otopathy [39], and developmental defects [40].

In the present study, we explored the molecular and cellular consequences of KCNQ4 deletion in the OC using a DFNA2-related model of HL. Our findings revealed significant disruptions in the distribution of key proteins essential for HC function, including abnormal localization of prestin and BK channels in OHCs, along with structural changes in their stereocilia. Both OHCs and SCs underwent apoptosis, while IHCs showed signs of degeneration, including abnormalities in ribbon synapses. Additionally, SGNs and satellite cells underwent apoptosis in aged KO mice. Notably, exposure to high levels of noise did not exacerbate HC damage in these mice.

## 2. MATERIALS AND METHODS

### 2.1 Animals

transgenic C3H/HeJ mice, which do not express the KCNQ4 protein (*Kcnq4*^−/−^ or KO) due to a deletion covering exons 6 through 8, were used in the study [6, 10]. Wild-type (WT) C3H/HeJ littermates (*Kcnq4*^+/+^) served as control animals and for comparison between strains. Mice of both sexes were included in all experiments to ensure that any observed effects were not sex-specific. The age categories were as follows: a) young mice, aged 4 to 20 postnatal weeks (W); b) middle-aged adults, aged 40 to 60 W; and c) older mice, aged over 60 W. The experimental procedures adhered to the ethical standards set by the Council for Care and Use of Experimental Animals (CICUAE, protocol no. 083/2016) of the Universidad Nacional del Sur (UNS), which comply with the European Parliament and Council directives (2010/63/EU).

### 2.2 Tissue preparation

Mice were euthanized by CO_2_ exposure and inner ears were promptly removed from temporal bones. In order to monitor HC and SGN degeneration, cochleae were studied by immunofluorescence (IF) using two different approaches i) mounted as a whole or, ii) in thin tissue sections. Both started with tissue fixation by overnight submersion in 4 % paraformaldehyde, washed with 1X PBS, and decalcified using 8–10 % EDTA in 1X PBS for up to 7 days depending on animal age, on a rocking shaker at 4 °C. Only for the autophagy analysis by IF, cochlea samples were fixated with 100 % methanol.

### 2.3 Whole-mount cochlear preparations

The OC was dissected from decalcified cochleae following a procedure previously published [10]. This technique involves dividing the cochlea into three longitudinal sections: basal, middle, and apical turns, using fine scissors. Next, the vestibular system, spiral ligament, modiolus, and tectorial membrane were removed, and the OC was carefully isolated. Lastly, the hook was separated from the remaining basal segment.

### 2.4 Cochlear sections

Whole inner ears were processed following protocols similar to those outlined by Spitzmaul et al. and Barclay et al. [41, 42]. In brief, after decalcification, the inner ears were cryoprotected by immersion in 15 % sucrose for 4 h, followed by 30 % sucrose overnight. The tissues were then embedded in OCT. Longitudinal 10-15 μm sections, aligned with the modiolus, were cut using a cryostat (Leica CM 1860) and stored at −20 °C until further processing.

### 2.5 Immunofluorescence in whole-mount cochlear preparations

We followed two different protocols due to the primary antibodies’ requirements. *Protocol 1:* Cochlear turns were postfixed in 4 % PFA during 20 min, washed three times in 1X PBS and incubated for 2 h in blocking solution (2 % BSA, 0.5 % Nonidet P-40 in 1X PBS). Primary antibodies were incubated for 48 h in carrier solution (PBS containing 1 % BSA, and 0.25 % Nonidet P-40). Subsequently, tissue was rinsed three times in 1X PBS. Secondary antibodies, diluted in the carrier solution, were incubated for 2 h at room temperature. After that, samples were washed three times in 1X PBS. Finally, cochlear turns were mounted unflattened in Fluoromount-G (Southern Biotech). The following primary antibodies were used: goat anti-prestin (1:400, cat#sc-22,692, Santa Cruz Biotechnology), rabbit anti-cleaved-caspase-3 (cCAS-3) (1:400, cat#9661, Cell Signaling), rabbit anti-LC3B (1:200, cat2775S, Cell Signaling), mouse anti-Kir4.1 (1:100, sc-293252, Santa Cruz Biotechnology), mouse anti-MaxiKα (BK) (1:100, sc-374142, Santa Cruz Biotechnology).

For *Protocol 2* there were a few modifications: cochlear turns were permeabilized by freeze in 30 % sucrose and blocked in 5 % normal goat serum with 1 % TritonX-100 in 1X PBS. The following primary antibodies, diluted in blocking buffer and incubated for 16 h at 37° C were used: mouse anti-CtBP2 (1:200, IgG1, #612044, BD Transduction Labs), mouse anti-GluA2 (1:2000, IgG2a, #MAB397, Millipore).

The following fluorescently-labeled secondary antibodies were obtained from Molecular Probes and used diluted (1:500): Alexafluor488-conjugated donkey anti-rabbit (A21206, Invitrogen), Alexafluor555-conjugated donkey anti-mouse (A31570, Invitrogen), AlexaFluor 488-conjugated goat anti-mouse IgG2a (A21131, Invitrogen), AlexaFluor 568 conjugated goat anti-mouse IgG1 (A21124, Invitrogen) and donkey anti-goat 633 (A-21082, Invitrogen). Nuclei were stained with DAPI (1:1000).

### 2.6 Immunofluorescence on cochlear sections

Tissue sections mounted on slides were post-fixed with 2 % PFA for 20 min, washed with 1X PBS, and then blocked in the same buffer as described in section 2.3. The sections were incubated overnight with primary antibodies: rabbit anti-beta III tubulin (TUBB3, 1:1000, Covance #mrb-435p) and mouse anti-Kir4.1 (1:100, sc-293252, Santa Cruz Biotechnology), both diluted in carrier solution. Secondary antibodies, including AlexaFluor488-conjugated donkey anti-rabbit (A21206, Invitrogen) and AlexaFluor555-conjugated donkey anti-mouse (A31570, Invitrogen), were applied for 1 h.

### 2.7 Hair cell counting and cytocochleogram plotting

Labeled cells in whole-mount cochleae were imaged using an epifluorescence microscope (Nikon Eclipse E-600) with a CCD camera (Nikon K2E Apogee) and a confocal microscope (Zeiss LSM900 with Airyscan module II). Images were analyzed with Image J software, and cell counting was performed manually. HCs were identified based on their location within the OC. A cytocochleogram is a method used to map the number and distribution of HCs along the cochlear length [10, 43, 44]. It is created by plotting the number of OHCs or IHCs against the relative distance from the apex. To standardize the length of the OC, the cochlea was divided into 20 segments (S), each representing 5 % of the total cochlear length, with 100 % representing the full length. Each segment was labeled according to its percentage of the total length, starting from the apex (e.g., 0 % to 5 % = S5) to the base (S100). For the analysis of cCAS-3 signal, apoptotic cells were identified by the co-localization of prestin and cCAS-3 in OHCs or Kir4.1 and cCAS-3 in supporting cells of the OC. Apoptotic cells were evaluated in a cytocochleogram between S20 and S80, identifying the cCAS-3 signal in each S. In the case of acoustic trauma, the cCAS-3 signal was specifically evaluated in the first 5 % of the middle turn (S40 to S70), as this region is particularly affected by acoustic overexposure.

### 2.8 Synaptic counts on HCs

Confocal z-stacks (with a 0.5 µm step size) from the medial regions of whole-mount cochleae were acquired using a Zeiss LSM900 microscope with an Airyscan module II, equipped with a 63x oil-immersion lens (1.4x digital zoom). The image stacks were imported into Fiji software, where HCs were identified based on their CtBP2-stained nuclei. Typically, each stack included 10–20 IHCs. A custom Fiji plugin was used to automate the quantification of synaptic ribbons, glutamate receptor clusters, and the co-localization of synaptic puncta. Each channel was analyzed separately, generating maximum projections to count CtBP2 or GluA2 puncta. A composite image was created by merging the two channels, using the CtBP2-stained nuclei as a reference to define regions of interest (ROIs) corresponding to each IHC. The maximum projections of the individual channels were merged into a 32-bit image, which was subsequently thresholded and converted into binary images. Automated particle counting was then performed within each ROI. In addition, each stack was analyzed using the Fiji “3D Object Counter” plugin in order to measure the volume of each puncta. The threshold was manually adjusted on a per-sample basis to ensure that protein signals were clearly distinguishable from the background [45]. CtBP2 signal volume in OHC was also estimated using this procedure.

### 2.9 Analysis of OHC membrane protein localization

Confocal z-stacks with three channels (prestin, BK, and DAPI) from the medial regions of each whole-mount cochlea were captured using the previously described setup. For the analysis of prestin and BK, X-Y Maximum Intensity Projections (MIPs) were created for the inner OHC row in both WT and KO animals. Using Fiji software, the nuclear perimeter labeled with DAPI and the portion of the nuclear perimeter covered by prestin-labeled membrane were measured. A ratio of these measurements was then calculated to determine the fraction of nuclei covered by prestin.

### 2.10 Auditory brainstem responses (ABRs) and distortion product otoacoustic emissions (DPOAEs)

Mice were anesthetized with xylazine (10 mg/kg i.p.) and ketamine (100 mg/kg i.p.). DPOAEs at 2f1–f2(f2/f1=1.2, and the f2 level 10 dB lower than the f1 level), were recorded with a custom acoustic assembly consisting of two dynamic earphones used as sound sources (CDMG15008–03A; CUI) and an electret condenser microphone (FG-23329-PO7; Knowles) coupled to a probe tube to measure sound pressure near the eardrum. Stimuli were generated digitally, while resultant ear-canal sound pressure was amplified and digitally sampled at 4 µs intervals (24-bit DAQ boards, NI PXI-1031; National Instruments). Fast Fourier Transforms were computed and averaged over five consecutive waveform traces, and 2f1–f2 DPOAE amplitude and surrounding noise floor were extracted. Iso-response curves were interpolated from plots of amplitude versus sound level, performed in 10 dB-steps of f1 level. ‘Threshold’ was defined as the as the lowest f2 level in which the signal to noise floor ratio is >1 For measurement of ABRs, needle electrodes were inserted at vertex and pinna, with a ground near the tail. ABRs were evoked with 5-ms tone pips (0.5-ms rise-fall, cos2 onset, at 40/s). The response was amplified (10,000X), filtered (0.3–3 kHz), and averaged with an A-D board in a LabVIEW-driven data-acquisition system. Sound level was raised in 10 dB-steps from 10 dB below threshold to 80 dB SPL. At each level, 1,024 responses were averaged (with stimulus polarity alternated), using an ‘artifact reject’ whereby response waveforms were discarded when peak-to-peak amplitude exceeded 15 µV. Upon visual inspection of stacked waveforms, ‘threshold’ was defined as the lowest SPL level at which any wave could be detected, usually the level step just below that at which the response amplitude exceeded the noise floor (± 0.25 µV). For amplitude versus level functions, wave-I peak was identified by visual inspection at each sound level and the peak-to-peak amplitude computed. The amplitudes of ABR peaks 1 and 2 were calculated through offline analysis of the peak-to-peak amplitude of the stored waveforms.

### 2.11 Acoustic overexposure

Animals were exposed free-field in the same acoustic chamber used for cochlear function tests. Noise calibration to target SPL was performed immediately before each acoustic overexposure. Acoustic trauma consisted of a 2 h exposure to a 1–16 kHz noise presented at 110 dB sound pressure level (SPL).

### 2.12 RNA isolation and retrotranscription

Cochlear RNA was extracted from young mice. For each experiment, samples from three to four mice were pooled. In brief, immediately after cochlear excision, tissue was immersed in ice-cold 1X PBS and then, the vestibular region was removed to keep only the cochlea. Total RNA was extracted using the BioZol reagent (PB-L, RA0202) and chloroform/isopropanol protocol. cDNA was produced from 500 ng total RNA with M-MLV Reverse Transcriptase (PB-L, EA1301) using anchored oligo (dT)s following manufacturer’s indications. For the cochlear nucleus RNA, brain portions containing the cochlear nucleus (-5.02 mm to -

6.48 mm from bregma, according to the Paxinos Atlas [46] ( were pooled from two young animals. Then, the procedures of RNA isolation and retrotranscription were similar to the described above.

### 2.13 Quantitative PCR (qPCR)

it was carried out using the cDNA generated previously employing the Sso Advanced Universal SYBR Green Supermix (BioRad) in a Rotor-Gene 6000 real-time PCR cycler (Corbett Research). The primers used for different genes were: Bax, 5′-GATCCAAGACCAGGGTGGC-3′ (forward) and 5′-CTTCCAGATGGTGAGCGAGG-3′ (reverse); Bcl-2, 5′-GGGGATGACTTCTCTCGTCG-3′ (forward) and 5′-CATGACCCCACCGAACTCAA-3′ (reverse); HPRT 5′-GTTCTTTGCTGACCTGCTGGA-3′ (forward) and 5′-AATGATCAGTCAACGGGGGA-3′ (reverse); GAPDH 5′-GAGAAACCTGCCAAGTATGATGAC-3′ (forward) and 5′-CCCACTCTTCCACCTTCGAT-3′ (reverse). Data analysis was done applying the ΔΔCt method [10, 47] to obtain relative mRNA expression (RQ).

### 2.14 Scanning electron microscopy (SEM)

Basal cochlear segments were dissected from young and middle-aged WT and KO mice. The cochleae were fixed in 2.5 % glutaraldehyde for 2 h and subsequently decalcified as previously described. After fixation, samples were dehydrated using ethanol and then processed using a critical point dryer. Finally, the tissues were examined using a LEO-EVO 40 XVP-EDS Oxford X-Max 50 scanning electron microscope, provided by the Microscopy Facility at CONICET Bahía Blanca.

### 2.15 Statistical analysis

For single mean comparisons, Student’s t-test was applied (Fig. 2C, D and E; Fig. 3B and D; Fig 5B; Fig 6D; Fig. 8D; Fig. S2). For multiple comparisons, two-way ANOVA, followed by Tukeýs post-hoc test, was used (Fig. 1C and D; Fig. 8A and B). At least three independent experiments were used. Statistical analyses were performed using Graph Pad 8.3 Software. Statistical significance is represented with (*) when p < 0.0500; (**) when p < 0.0100; (***) when p < 0.0010 and (****) when p < 0.0001.

**Figure 1:**
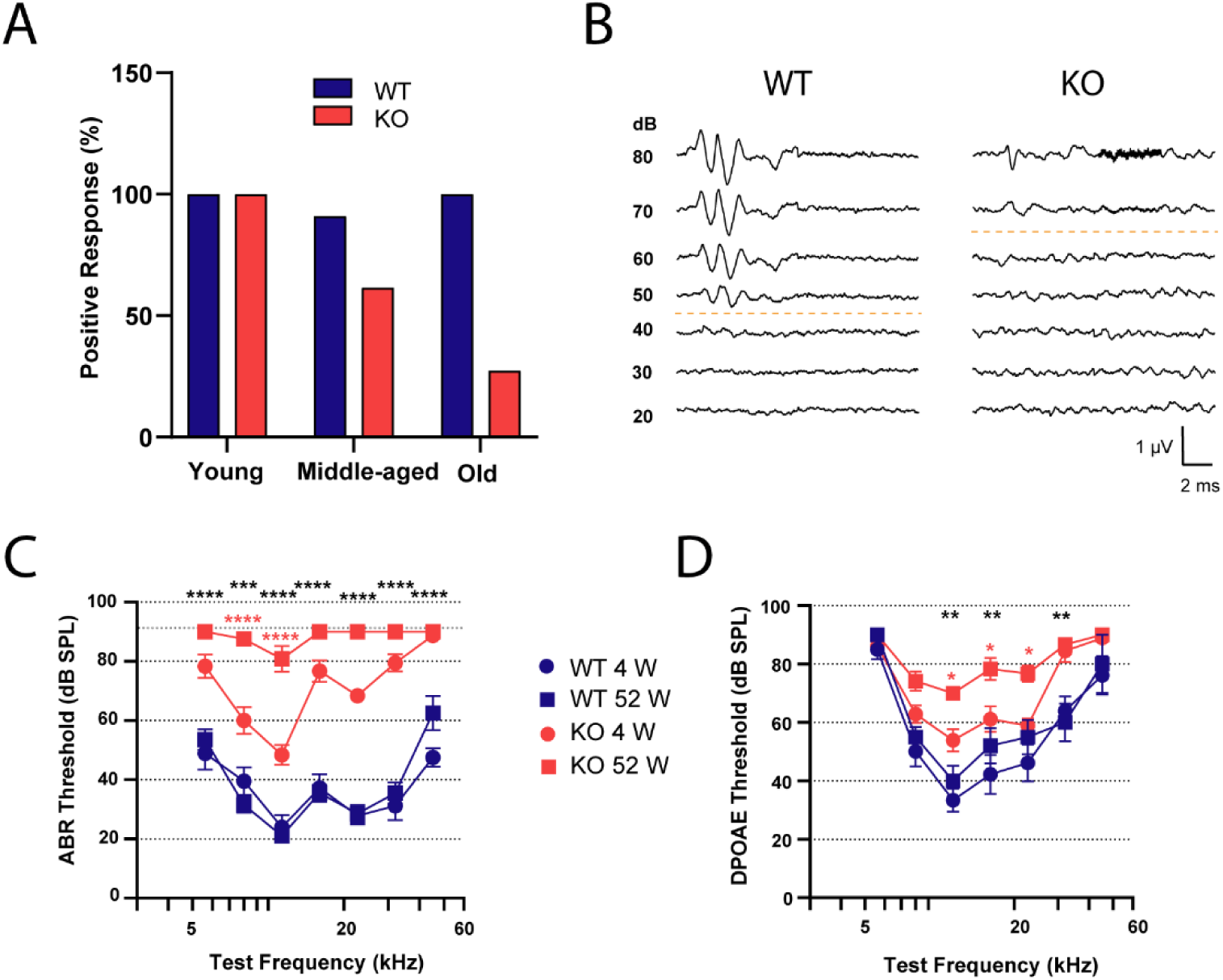
Comparative auditory function assessment in WT and KO mice across age groups. A. Preyeŕs reflex test. Graph depicting the percentage of WT (blue) and KO (red) animals at different ages that exhibited a positive response to the test (n = 11 - 22). **B. ABR recordings.** Representative traces of evoked potentials from WT and KO young animals at different sound intensities stimulated at 16.00 kHz. Dashed lines indicate the threshold used for quantification. **C, D. Auditory function measurements.** ABR (**C**) and DPOAE (**D**) thresholds for WT (blue) and KO (red) animals were determined at 4 W (circles) and 52 W (squares) across a frequency range of 5.60 - 45.25 kHz. Data are presented as mean ± SEM. Statistical analysis was performed using two-way ANOVA followed by Tukeýs post hoc test. *, p < 0.0500. **, p < 0.0100. ***, p < 0.0010. ****, p < 0.0001 (n = 9). Red asterisks indicate comparisons between age groups in KO animals, while black asterisks indicate comparisons between WT and KO animals at 4 W.

**Figure 2.**
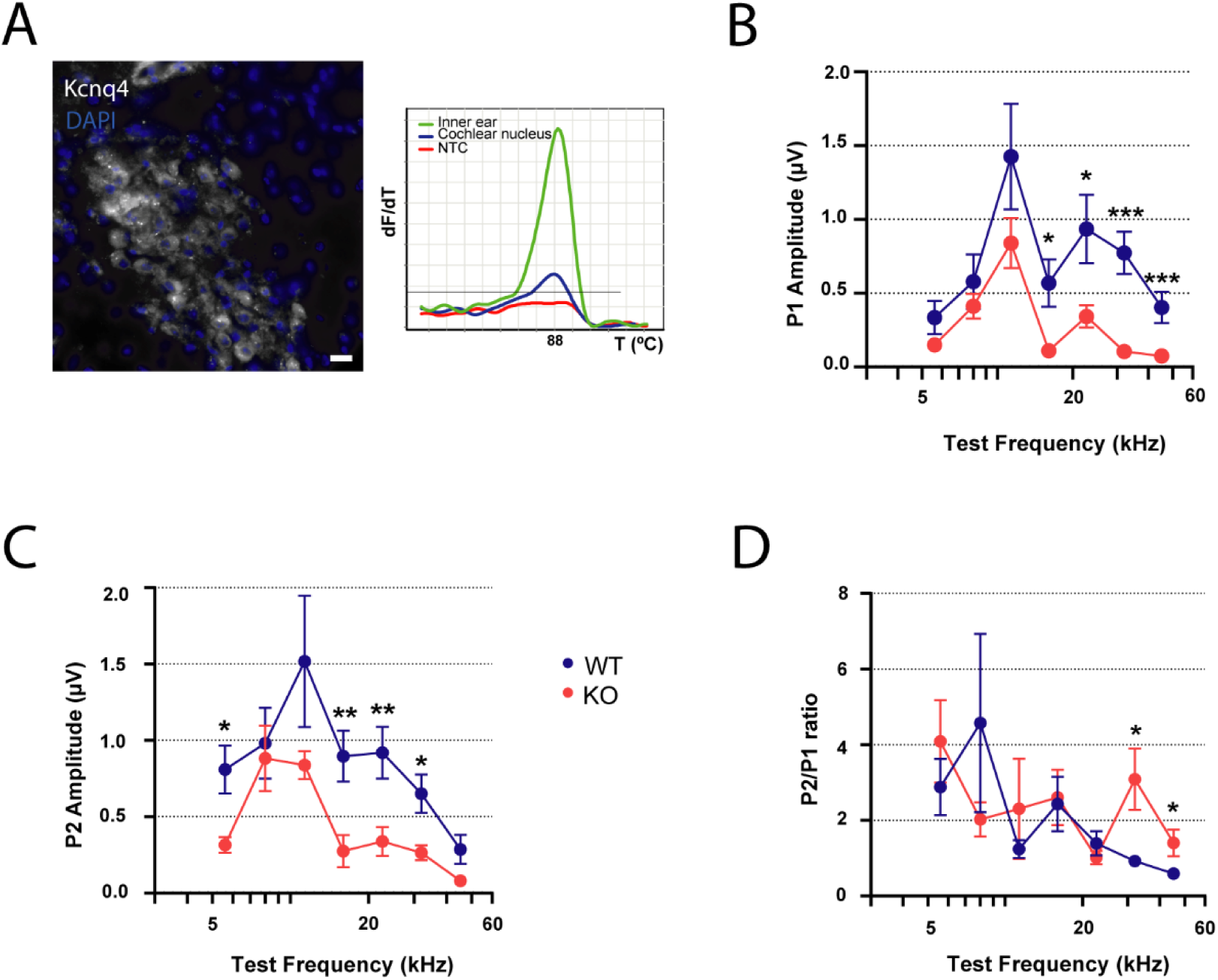
Suprathreshold ABR peaks analysis in KCNQ4 KO mice. A. KCNQ4 expression in cochlear nucleus. *Left*, KCNQ4 immunostaining (white) in the cochlear nucleus area of a middle-aged animal. Scale bar: 10 µm. *Right*, melting curve of *Kcnq4* qPCR product from cochlear nucleus samples (blue line) and inner ear samples (as a positive control, green line). **B, C. ABR peak amplitudes.** P1 (**B**) and P2 (**C**) amplitudes at 80 dB SPL in young WT (blue) and KO (red) animals across a frequency range of 5.60 - 45.25 kHz. Data are presented as mean ± SEM. Statistical analysis was performed using Student’s t-test at each frequency. *, p < 0.0500. **, p < 0.0100. ***, p < 0.0010 (n = 9). **D. P2/P1 ratio analysis.** Relationship between P2 and P1 values at all tested frequencies. Data are presented as mean ± SEM. Statistical analysis was performed using Student’s t test at each frequency. *, p < 0.0500 (n = 9).

**Figure 3:**
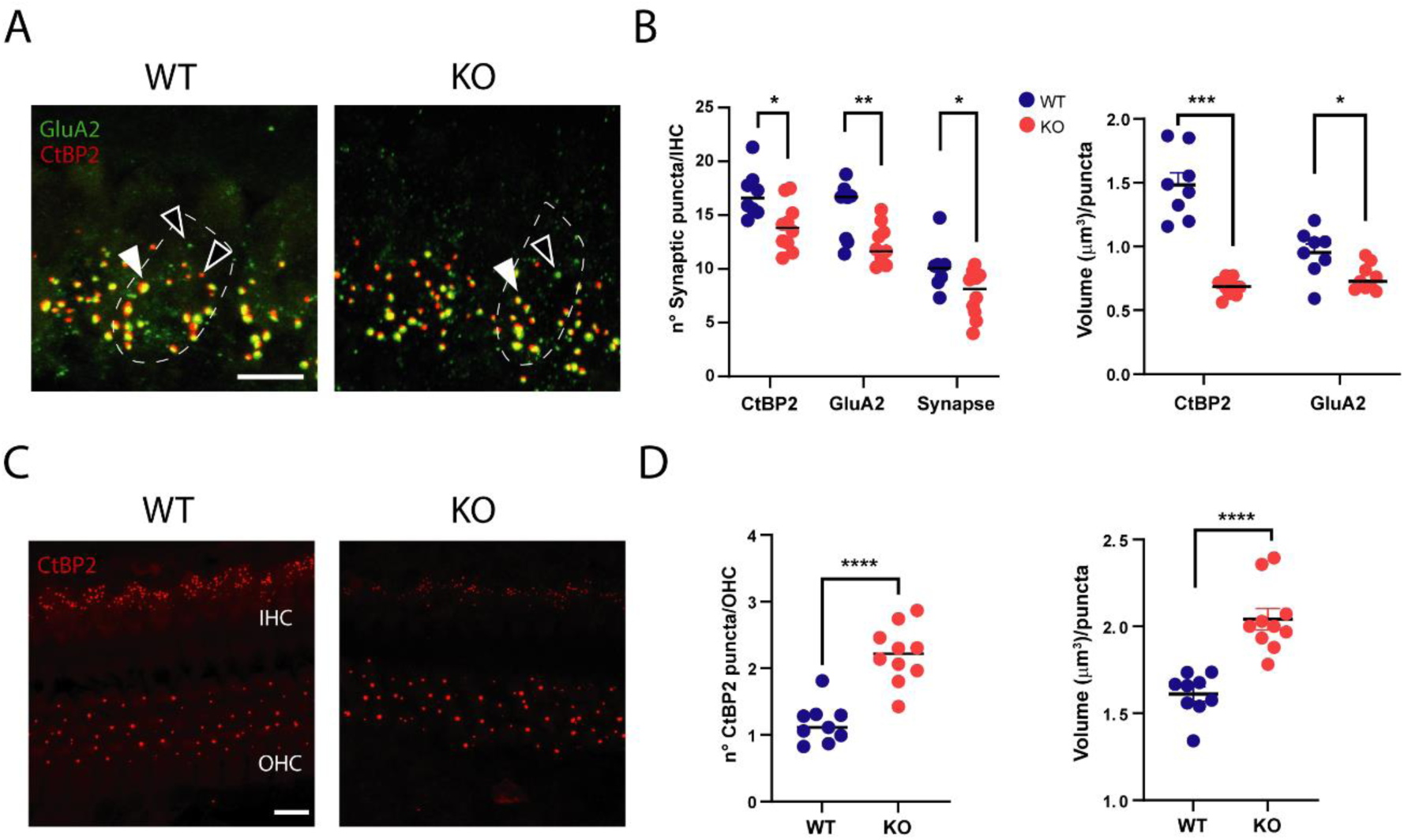
Impact of *Kcnq4* deletion on afferent synaptic contacts in inner and outer hair cells. A. Afferent synapses in IHCs. Confocal images showing CtBP2 (red) and GluA2 (green) staining in IHCs from both genotypes at 4 W. Dashed lines delineate individual IHCs. Filled arrowheads indicate afferent synapses while hollow arrowheads mark individual CtBP2 or GluA2 puncta. Scale bar: 10 µm. **B. Quantification of synaptic markers in IHCs.** *Left*, the number of CtBP2, GluA2, and afferent synapse signals per IHC from WT (blue) and KO (red) mice. *Right*, Volume of CtBP2 and GluA2 puncta in both genotypes. Data are presented as mean ± SEM (n = 8 - 10). Statistical analysis was performed using Student’s t-test. *, p < 0.0500; **, p < 0.0100; ***, p < 0.0010. **C. Afferent synapses in OHCs.** Confocal images of CtBP2 (red) staining in OHCs from 4 W animals of both genotypes. Scale bar: 10 µm. **D. Quantification of CtBP2 signals in OHCs.** Number of CtBP2 signals per OHC (left) and volume of CtBP2 puncta (right) from WT (blue) and KO (red) animals. Data are presented as mean ± SEM (n = 10). Statistical analysis was performed using Student’s t-test. ****, p < 0.0001.

**Figure 4:**
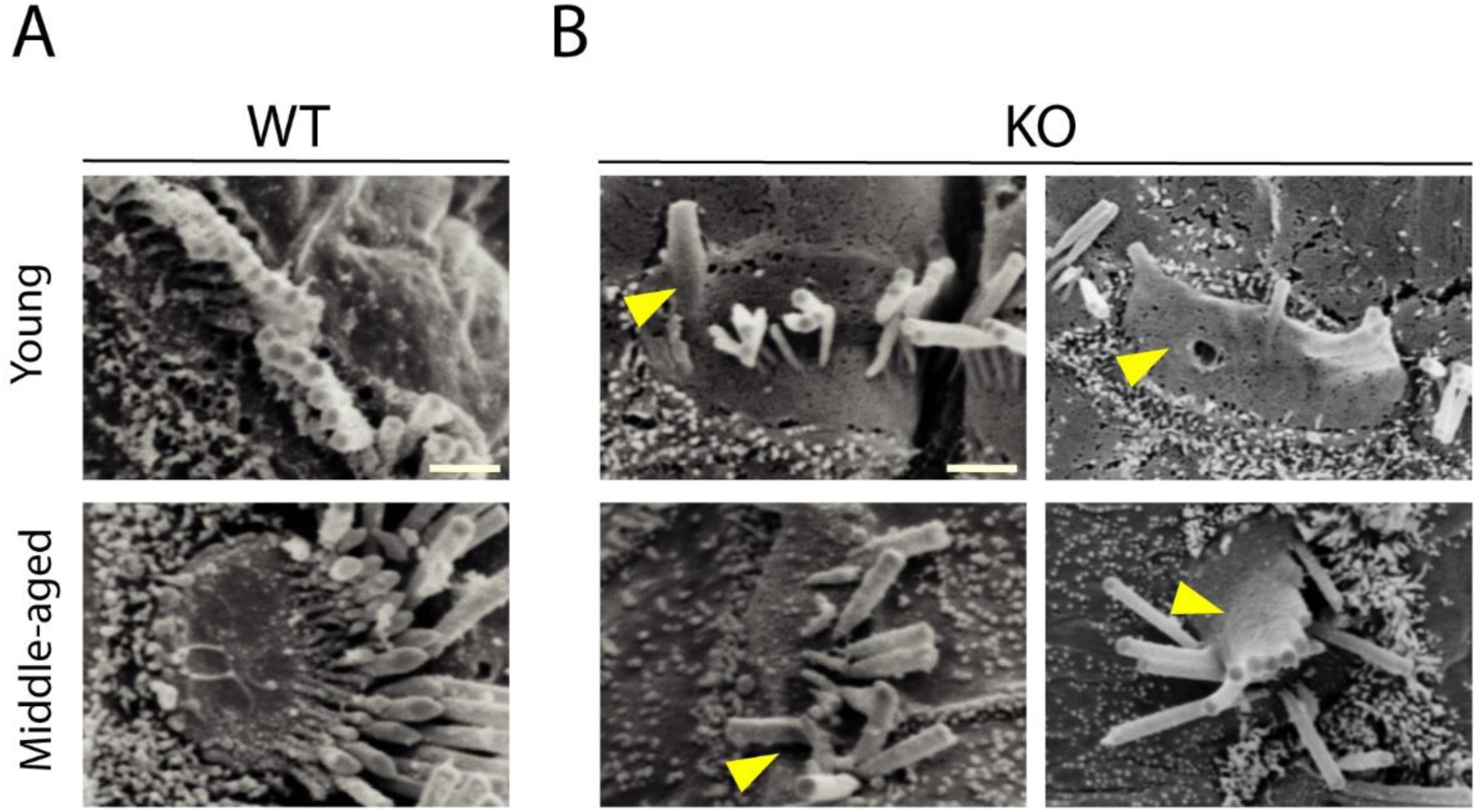
Ultrastructural alterations in the apical surface of inner hair cells in KO Mice. Ultrastructural analysis of hair bundles in IHCs using scanning electron microscopy (SEM). Representative images from WT (**A**) and KO mice (**B**) are shown. Pictures correspond to young and middle-aged animals of both genotypes. Yellow arrowheads indicate different structural alterations such as fused stereocilia, floppy stereocilia, among others. Total magnification: 30000X. Scale bar: 1 µm.

**Figure 5.**
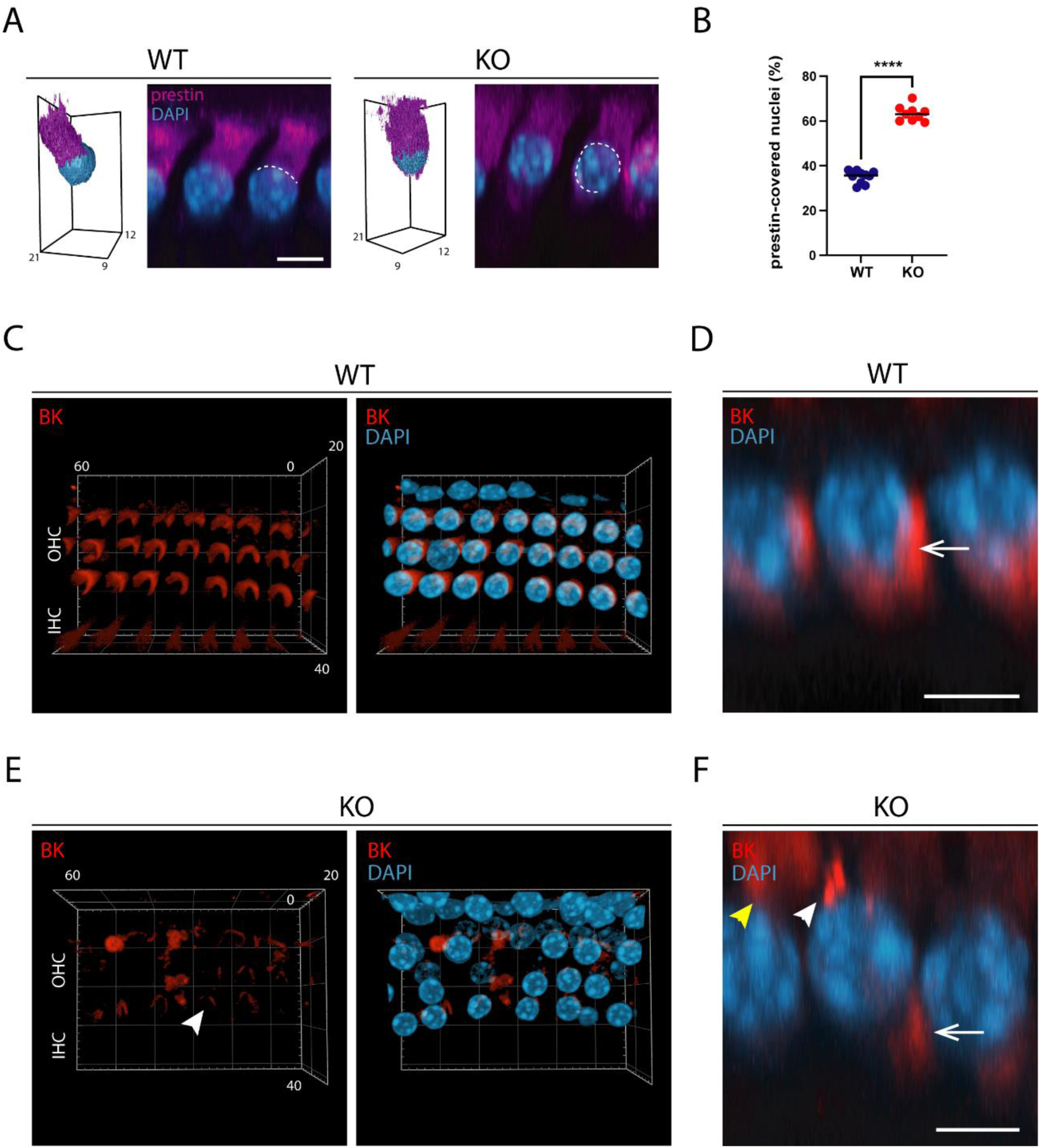
Effect of *Kcnq4* deletion on the localization of hearing-related membrane proteins in OHCs. **A.** Prestin localization in OHCs from young WT and KO animals. On the l*eft* of each genotype, 3D images depict prestin (violet) and DAPI (blue) staining in individual OHCs. The 3D scale (µm) is indicated in the image. On the r*ight* of each genotype, confocal XY-plane MIP images show prestin (violet) and DAPI (blue) staining in a single row of OHCs. Dashed lines indicate prestin-covered surface of the nuclei. Scale bar: 5 µm. **B. Quantification of prestin-covered nuclei.** Bar graph showing the percentage of nuclear surface covered by prestin in WT (blue) and KO (red) mice. Data are presented as mean ± SEM. Student’s t-test. ****, p < 0.0001 (n = 9). **C, E. BK expression in young animals.** 3D confocal images showing BK (red) and DAPI (blue) staining in OHCs from WT (**C**) and KO (**E**) mice. The 3D scale (µm) is indicated in the image. **D, F. Lateral view of BK expression in OHCs.** XY-plane MIP pictures of BK (red) and DAPI (blue) staining in OHCs from WT (**D**) and KO (**F**) mice. White arrow indicates BK localization in the subnuclear region. White arrowhead indicates a compact signal of BK in the supranuclear region, and yellow arrowhead indicates diffuse signals in the same region. Scale bar: 5 µm.

**Figure 6.**
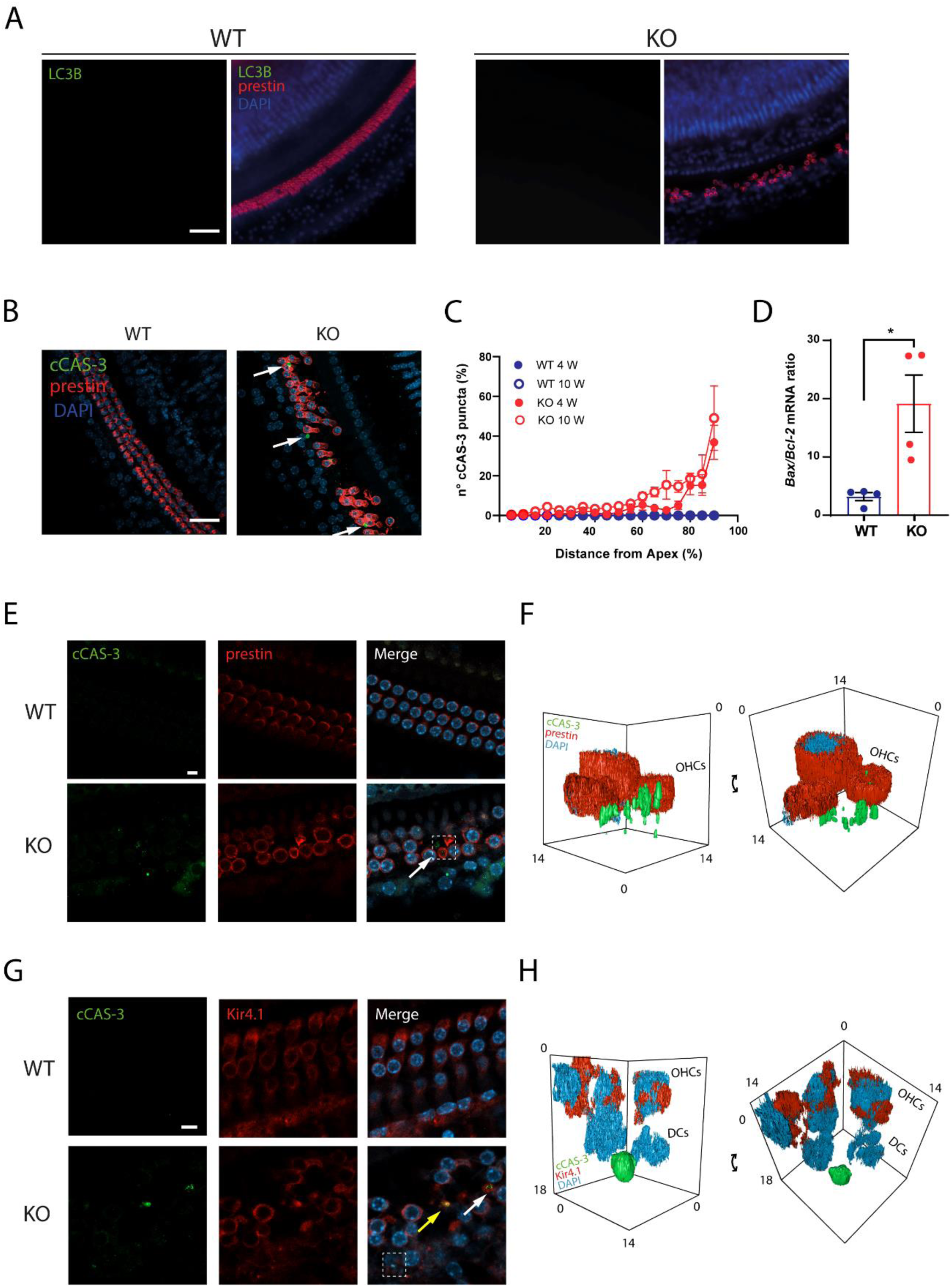
Mechanism of cell death in KCNQ4 KO mice. **A. Autophagy evaluation in young animals**. LC3B (green), prestin (red) and DAPI (blue) staining in WT and KO animals of 4 W. Scale bar: 10 µm. **B. cCAS-3 staining in organ of Corti.** *Left*, images from WT and KO animals of 10 W, showing prestin (red), cCAS-3 (green) and DAPI (blue) staining. White arrows indicate cCAS-3 signal. Scale bar: 30 µm. **C.** Quantification of cCAS-3 signal expressed in percentage of number of cCAS-3 puncta in 4 W WT (blue filled circles), 10 W WT (blue hollow circles), 4 W KO (red filled circles) and 4 W KO (red hollow circles). (n = 4 - 6). **D. *Bax*/*Bcl*-2 mRNA ratio.** The ratio between *Bax* and *Bcl*-2 mRNA was evaluated by qPCR. Four cochleae-pooled samples of 3 animals each were used. Data are presented as mean ± SEM. Student’s t-test. *, p < 0.0500. **E. Prestin and cCAS-3 staining in OHCs**. Confocal microscopy images showing cCAS-3 (green), prestin (red) and DAPI (blue) staining in WT and KO of 4 W animals. White arrow indicates cCAS-3 signal. Scale bar: 5 µm. **F.** 3D surface images from white dashed box indicated in panel E. Z-angled front view (right). 3D scale set in image. **G. Kir4.1 and cCAS-3 staining in OHCs**. Confocal images showing cCAS-3 (green), Kir4.1 (red) and DAPI (blue) in WT and KO of 4 W animals. White arrow indicates cCAS-3 signal and yellow arrow indicates co-localization of cCAS-3 signal with Kir4.1 signal. Scale bar: 5 µm. **H.** 3D surface images from white dashed box indicated in panel G. Z-angled front view (right). Scale set in image. DCs: Deiter’s cells.

## 3. RESULTS

### 3.1. Functional Studies

KO mice exhibited HCs loss at different time points [10]. OHCs begin to die as early as 3 W in the basal turn of the mouse cochlea while IHCs and SGNs are lost after 40 W in both the basal and apical turn. Although all HCs express the KCNQ4 channel and, in its absence, all HCs are depolarized by the time of hearing onset [6], cell loss follows a strict pattern similar to that observed in ARHL and/or NIHL [3, 11, 48]. Before HC loss, hearing function is impaired, but the magnitude of this process in young and middle-aged animals was not determined. To explore auditory function in KO mice, we performed different functional studies.

#### 3.1.1: Preyer Test

This semiquantitative test assesses the automatic (unconscious) startle response of animals to a loud sound, set at 115 dB SPL in this study. It serves as a quick screening tool for detecting profound deafness. Results showed that nearly 100 % of WT mice across all age groups showed a positive response (20/20 for young, 18/20 for middle-age, and 14/14 for aged animals; Fig. 1A). In KO animals, however, the positive response rate declined significantly with age: 100 % in young animals (18/18), 60 % in middle-aged (8/13), and 30 % in aged animals (3/11) (Fig. 1A).

#### 3.1.2: Auditory Brainstem Response

To evaluate cochlear functionality, we analysed sound-evoked potentials along the ascending auditory pathway in both WT and KO mice. We first determined the auditory sensitivity across cochlear turns in young and middle-aged mice by measuring ABRs thresholds at frequencies spanning from 5.60 kHz to 45.25 kHz. The auditory threshold was determined as the first identifiable peak signal above background response, occurring approximately 1-2 ms after the auditory stimulus. As shown in Fig. 1B, at a sound frequency of 16 kHz, the threshold for young WT mice was around 40 dB, while in young KO animals, this value was notably elevated.

Auditory thresholds for both genotypes displayed a U-shaped pattern, indicating greater auditory sensitivity in the middle cochlear region compared to the boundaries of the frequency range (Fig. 1C). In WT animals at both ages, this typical pattern for the C3H/HeJ mice strain [48, 49], showed threshold of around 20-40 dB within the 8.00 to 32.00 kHz range, and around 50 dB at the range boundaries (Fig 1C, blue symbols). In contrast, young KO mice exhibited significantly higher auditory thresholds across all frequencies (p = 0.0003 for 8 kHz and p < 0.0001 for the others, two-way ANOVA, Tukeýs post hoc test). However, the shift was not uniform across the range. Threshold shifts were 30, 20 and 20 dB at 5.60, 8.00 and 11.33 kHz, respectively, and exceeded 40 dB at higher frequencies (Fig. 1C, red circles).

At 52 W, WT mice displayed ABR thresholds similar to those observed at 4 W (Fig. 1C, blue squares vs. blue circles), indicating that auditory function was well-preserved in the C3H/HeJ strain throughout aging. In contrast, middle-aged KO animals exhibited higher auditory threshold than young KO animals, exceeding the upper intensity value of 90 dB SPL for 5.60, 16.00, 22.65, 32.00 and 45.25 kHz (Fig. 1C, red squares vs red circles). For 8.00 and 11.33 kHz frequencies, statistically significant differences were determined (Fig. 1C, red symbols). In consequence, comparing 52 W WT vs KO animals we found a significant increase in auditory threshold at all frequencies, assuming an upper limit value of 90 dB (p < 0.0001, two-way ANOVA, Tukeýs post hoc test).

These observations collectively show that while auditory function remains well-preserved in middle-aged WT mice, KO have a progression of mild to severe auditory impairment.

#### 3.1.3: Distortion Product of Otoacoustic Emissions

To evaluate OHC function, we measured DPOAE across the frequency range of 5.60 to 45.25 kHz in both young and middle-aged WT and KO mice. In WT mice, DPOAE thresholds at 4 W showed a region of high sensitivity within the 8.00-22.65 kHz range, with values around 30-50 dB SPL. Sensitivity was reduced at the boundary frequencies, with higher thresholds observed at both the lowest (5.60 kHz) and highest (32.00 and 45.25 kHz) ends of the range (Fig. 1D, blue circles). At the same age, KO animals showed thresholds of ∼20 dB higher than WT, although statistical differences were observed only for 3 frequencies (p < 0.0100 for 11.33, 16.00 and 32.00 kHz; two-way ANOVA, Tukeýs post hoc test) (Fig. 1D, red vs blue circles).

In middle-age WT mice, DPOAE thresholds remained similar to those measured at 4 W (Fig. 1D) (p > 0.0500 for all of them, two-way ANOVA (not indicated in the graph)). Middle-aged KO animals showed a threshold increase compared to young KO mice in the range from 11.33 to 22.65 kHz of around 20 dB (Fig. 1D) (p < 0.0500 for all of them, two-way ANOVA, Tukeýs post hoc test (red asterisks)). Comparing WT vs KO animals at this age, threshold values further increased compared to young conditions. Differences in the threshold shift were detected at 5 frequencies (Fig. 1D) (p < 0.0500 for 8.00 kHz and 22.65 kHz; p < 0.0100 for 16.00 kHz and 32.00 kHz; p < 0.0010 for 11.33 kHz, two-way ANOVA, Tukeýs post hoc test (not indicated in graph)). These elevations suggest that OHC amplification is reduced since 4 W in the KO mice in the middle cochlear length.

#### 3.1.4: Suprathreshold analysis

To investigate if auditory processing to the brainstem is altered, we performed suprathreshold analysis of peak waves obtained from ABR traces. As KCNQ4 expression has been reported in cochlear nucleus (CN) in the auditory pathway, we first performed IF staining of midbrain regions to confirm KCNQ4 expression in CN of C3H/HeJ mouse strain. As already published [7], we observed labeling of KCNQ4 in neurons from the ventral CN (Fig. 2A). To confirm the presence of KCNQ4 by a complementary technique to IF, we analyzed the expression of its gene, *Kcnq4*, by qPCR, using cochlear tissue as a positive control for gene expression (Fig. 2A, right). The melting curve at 87.5°C corresponds to the *Kcnq4* product, while no amplification was detected in the negative control (absence of sample). Because KCNQ4 is expressed in IHCs and the CN, where it plays a key role in auditory processing to the CNS, we focused our analysis on peaks 1 and 2 of the ABR recordings. The ABR growth function from 20 to 80 dB SPL and the peak latency of the measurements were analyzed for each frequency (Suppl. Fig. 1 and Table 1). In WT mice, P1 increases linearly with the sound intensity for most of the frequencies (Suppl. Fig 1). In KO animals, this behavior is more dependent on the stimulation frequency. Due to the increased hearing threshold, there is a shift in the onset of P1 detection from 5.65 to 11.33 kHz. In this frequency range, once this peak is detected, there are practically no significant differences compared to the WT values. Meanwhile, from 16.00 to 45.25 kHz, P1 was lower for all sound intensities in KO mice compared to WT animals (Suppl. Fig. 1). A similar behavior was determined for P2. WT animals exhibited a linear increase with sound intensity at all frequencies while KO animals showed a similar pattern as observed for P1 at the same frequencies (Suppl. Fig. 1).

**Table 1.** ABR peak latency quantification at 4 W.

|  | Latency to Peak I |  | Latency to Peak II |  | Latency to Peak III |  | Latency to Peak IV |  | Latency to Peak V |  |
| --- | --- | --- | --- | --- | --- | --- | --- | --- | --- | --- |
|  | WT | KO | WT | KO | WT | KO | WT | KO | WT | KO |
| **5.60** | 1.64 ± | 1.77 ± | 2.59 ± | 2.72 ± | 3.42 ± | 3.63 ± | 4.77 ± | 4.65 ± | 6.02 ± | 5.92 ± |
| **kHz** | 0.12 | 0.22 | 0.13 | 0.12 | 0.41 | 0.22 | 0.38 | 0.28 | 0.75 | 0.26 |
| **8.00** | 1.51 ± | 1.67 ± | 2.40 ± | 2.45 ± | 3.26 ± | 3.37 ± | 4.47 ± | 4.52 ± | 5.61 ± | 5.48 ± |
| **kHz** | 0.15 | 0.17 | 0.22 | 0.24 | 0.38 | 0.36 | 0.56 | 0.22 | 0.72 | 0.34 |
| **11.33** | 1.46 ± | 1.51 ± | 2.28 ± | 2.41 ± | 3.22 ± | 3.20 ± | 4.35 ± | 4.25 ± | 5.36 ± | 5.29 ± |
| **kHz** | 0.15 | 0.24 | 0.32 | 0.41 | 0.39 | 0.39 | 0.53 | 0.39 | 0.55 | 0.23 |
| **16.00** | 1.47 ± | 1.54 ± | 2.25 ± | 2.33 ± | 3.17 ± | 3.07 ± | 4.24 ± | 3.89 ± | 5.33 ± | 5.01 ± |
| **kHz** | 0.17 | 0.23 | 0.29 | 0.42 | 0.48 | 0.47 | 0.55 | 0.49 | 0.46 | 0.59 |
| **22.65** | 1.31 ± | 1.44 ± | 2.21 ± | 2.17 ± | 3.09 ± | 3.16 ± | 4.10 ± | 3.85 ± | 5.13 ± | 4.77 ± |
| **kHz** | 1.37 | 0.11 | 0.30 | 0.18 | 0.47 | 0.44 | 0.55 | 0.50 | 0.55 | 0.58 |
| **32.00** | 1.30 ± | 1.46 ± | 2.14 ± | 2.10 ± | 2.95 ± | 3.04 ± | 4.00 ± | 3.83 ± | 5.06 ± | 4.84 ± |
| **kHz** | 0.14 | 0.16 | 0.28 | 0.23 | 0.42 | 0.49 | 0.51 | 0.48 | 0.42 | 0.48 |
| **45.25** | 1.23 ± | 1.23 ± | 1.93 ± | 2.01 ± | 2.55 ± | 2.84 ± | 3.42 ± | 3.52 ± | 4.48 ± | 4.18 ± |
| **kHz** | 0.25 | 0.17 | 0.28 | 0.20 | 0.51 | 0.48 | 0.56 | 0.52 | 0.85 | 0.45 |

At 4 W, P1 amplitude for WT mice increased with frequency, peaking at 11.33 kHz, and then gradually decreased up to 45.25 kHz (Fig. 2B). In KO animals, P1 amplitudes were reduced, with significant differences observed from 16.00 kHz onwards at 80 dB sound intensity (Fig. 2B) (p < 0.0500 at 16.00 kHz and 22.65 kHz; p < 0.0001 for 32.00 kHz and 45.25 kHz; two-way ANOVA, Tukeýs post hoc test). A similar trend was obtained for P2 amplitudes, which were consistently lower in KO mice. However, signal decrease was also significant at 5.65 kHz (Fig. 2C). To assess for central gain compensation in CN, we determined the ratio between P1 and P2 amplitude. As shown in Fig. 2D, the P2/P1 ratios were similar at low and middle frequencies but significantly higher for KO animals at 32.00 and 45.25 kHz.

Finally, we analyzed peak latency, a measure of neural transmission efficiency along the auditory pathway [50]. No differences in latency were observed between WT and KO mice for any peak (Table 1).

### 3.2. Structural changes in HCs

#### 3.2.1 Analysis of Afferent Synapses on Hair Cells

Our findings indicate that, in addition to the anticipated OHC dysfunction contributing to elevated ABR thresholds in KO mice, other potential alterations may exist. These could include impairments in IHCs, neurons of the auditory pathway, or the synaptic connections between IHCs and auditory nerve fibers. To assess this, we examined the synapses between IHC and auditory fibers through IF using antibodies targeting the presynaptic ribbon protein CtBP2-Ribeye and the postsynaptic GluA2 AMPA-type glutamate receptor on whole-mount tissues of the OC. Individual signals for each protein were imaged using confocal z-stack and quantified (Fig. 3A and B). The colocalization of puncta signals for CtBP2/GluA2 on MIP z-stack projections from confocal images was quantified as IHC-afferent synapses (Fig. 3A, white arrowheads). As shown in Fig. 3A, both genotypes exhibited the presence of CtBP2 and GluA2 in the IHC-afferent fiber synapse region. We counted the number of CtBP2 puncta in 6-12 IHCs per mouse in young animals and found a significant decrease in the number of this protein in the OC from KO mice (Fig. 3B, left) (p < 0.0500, Student’s t-test). The same trend was determined for the postsynaptic GluA2 signals (Fig. 3B, left) (p < 0.0100, Student’s t-test). Most of CtBP2 and GluA2 signals colocalized on MIP pictures (Fig. 3A, white arrowheads), however, some minor proportions were observed alone (Fig. 3A, hollow arrowheads). The number of synapses showed a significant decrease in KO animals (Fig. 3B, left) (p < 0.0500, Student’s t-test). We also determined the volume of each protein signal and found a decrease for both in KO mice (p < 0.0001 and p < 0.0500 for CtBP2 and GluA2, respectively, Student’s t-test). The most significant decrease was observed in the volume of presynaptic ribbons, which was approximately 2.2-folds smaller than in WT mice (Fig. 3B, right). Ribbon-like structures are also present in OHCs (Fig. 3C) [51]. In accordance with other publications [52] we found that there are around 1.1 ribbons per OHC in WT mice (Fig. 3D). This value increased by 1.9-fold in KO animals (Fig. 3D, left) (p < 0.0001, Student’s t-test). Additionally, we analyzed the volume of CtBP2 puncta in OHCs and observed a 1.3-fold increase in ribbon volume in KO animals (Fig. 3D, right) (p < 0.0001, Student’s t-test). Conversely, GluA2 signals were undetectable in both WT and KO in the synaptic are of the OHCs (not shown).

#### 3.2.2 Electron microscopy on apical surface of IHCs

Another possibility that explains ABR changes in KO animals could be the alteration in IHĆs hair bundle. In a previous work, we determined that the OHCs showed structural alterations in their hair bundles in young KO mice [10]. Now we analyzed the structure of hair bundles in IHCs by electron microscopy (Fig. 4). IHCs from young WT mice showed the typical distribution of stereocilia in 3-4 staircase rows for each HC bundle (Fig. 4A) [53, 54]. Although no loss of IHCs was observed at this age [10], KO animals exhibited several alterations in the hair bundle structure. First, we observed a decrease in the number of stereocilia and detected the loss of 1 or 2 rows of stereocilia located on the staircase of the hair bundle (Fig. 4B). Figure 4B (left) shows elongation and floppy stereocilia in some of them while on the right (top), the whole bundle was completely fused forming a unique structure. In middle-aged animals, when IHCs are degenerating, we observed more structural alterations in hair bundle from KO mice in the middle turn, such as basal and lateral fusion of two or more stereocilia, elongation of stereocilium and floppy ones (Fig. 4B, bottom).

#### 3.2.3 Analysis of membrane compartmentalization in OHCs

ABR threshold shift in apical and middle cochlear regions in young KO animals, where OHCs are well preserved, suggests an alteration in their function. The proper amplification of sound signal relies on the accurate localization of essential proteins including prestin, KCNQ4 and BK, among others. To evaluate this, we first determined the membrane localization of prestin in young mice. In WT animals, prestin is mainly localized in the lateral domain, from the uppermost part of the nuclear region up to the beginning of the apical domain (Fig. 5A, left). In KO mice, prestin localized not only in the lateral domain but also extends irregularly into the basal domain (Fig. 5A, right). To quantify the extent of prestin mislocalization, we calculated the proportion of the nuclear perimeter labeled with DAPI shielded by prestin-labeled membrane (Fig. 5A, white dashed line). We found a low percentage of the cell nucleus covered by prestin in WT animals (approx. 30 %), while it greatly increases in OHCs from KO mice to an average of 65 % (Fig. 5B) (p < 0.0001, Student’s t-test).

In the second step, we analyzed the localization of the BK channel in OHCs. In WT animals, BK signal formed well-defined uniform patches. From an upper view, they exhibited a C-shape pattern, oriented toward the lateral wall (facing the stria vascularis) (Fig. 5C). In a lateral view, we determined that the BK signal is located at the base of the OHC and covering the lower part of the nucleus (Fig. 5D, white arrow). In the KO mouse, BK signal, in an upper view, exhibited an irregular C-shape pattern with weaker signal intensity compared to WT (Fig. 5E). In the lateral view of the 3D-z-stack image, BK signal showed discrete patches in the basal region (Fig. 5F, white arrow) but also broader lateral membrane areas with weaker intensity (Fig. 5F yellow arrowhead) and compact areas with high signal intensity (Fig. 5F, white arrowhead).

In summary, our results indicate that both HCs present structural alterations that would impact in their transduction function.

### 3.3. Mechanism of Cell Death

The mechanism by which HCs are lost is unknown. For this reason, we analyzed mechanisms that may lead to cell death.

One of the mechanisms implicated in HL is autophagy, which can occur due to the accumulation of misfolded proteins or increased oxidative stress [55–58]. Besides, OHCs from KO animals exhibited accumulation of cytoplasmic vacuoles [6] which might be a result of increased autophagy [55]. We evaluated this mechanism by assessing the expression of the LC3B protein signal using IF. To verify antibody reactivity, we performed a staining in cell cultures under stress conditions, such as starvation or generation of oxidative stress (in the presence of 150 µM hydrogen peroxide). In both cases, we detected LC3B cytoplasmic signals adjacent to the cell nucleus, which were not present under normal conditions (Suppl. Fig. S2). We examined LC3B expression in cochlear whole-mount preparations from young and middle-aged mice in all cochlear turns. However, we did not detect LC3B signal in either HCs or SGNs in WT and KO mouse cochlear tissues at either age (Fig. 6A).

Another mechanism associated with cell death and HL is apoptosis [59]. We subsequently analyzed this mechanism in HCs and SGNs in both young and middle-aged animals of both genotypes. We evaluated the expression of the late apoptotic effector cCAS-3 using IF. For young WT animals, cCAS-3 signal was absent in all cochlear turns. In KO mice cCAS-3 signal was present in the OHC area in the middle and basal turns while it could not be detected in IHCs (Fig. 6B). To measure the proportion of cell in apoptosis, we counted the number of cCAS-3 signals alongside the entire cochlea. Quantitative analysis from KO mice revealed that the basal turn exhibited the highest amount of cCAS-3 expression and it increases at 10 W gradually towards the apical turn only in OHCs area (Fig. 6C). Additionally, to evaluate the degree of apoptotic activation, we evaluated the gene expression of pro- and anti-apoptotic factors *Bax* and *Bcl-2*, respectively, in cochlear tissues for both genotypes. We calculated the *Bax*/*Bcl-2* ratio for WT and KO mice. WT ratio value was 4.80 meanwhile for KO it increased to 39.79 (Fig. 6D) (p < 0.0010, Student’s t-test). In consequence, the expression ratio increased approximately 8-folds. Our results indicate an activation of the intrinsic apoptotic pathways that would be triggered during OHC degeneration in early stages.

We next analyzed which cell type from KO animals exhibited apoptotic markers by cCAS-3 expression using two cell population markers: prestin for OHCs and Kir4.1 for SCs. Figure 6E shows that young KO animals exhibited cCAS-3 signal in the OHC area. Some of them are intracellular, surrounded by membrane prestin signal and others are in empty spaces where OHCs were missing. A 3D magnification of the cCAS-3 signal in two different angles confirms its localization in the OHC region, indicating that it may corresponds to the final steps of apoptotic OHCs (Fig. 6F).

On the other hand, some of the cCAS-3 signals were observed outside the OHC region in the OC (Fig. 6E and G). We analyzed if this signal was generated by apoptotic SCs using Kir4.1 as a cell marker. cCAS-3 staining was present intracellularly in some SCs (surrounded by Kir4.1) (Fig. 6G, white arrows) while in others it colocalizes with the marker (Fig. 6G, yellow arrow) and in other cases, cCAS-3 was present in the absence of Kir4.1 signal (Fig. 6G, white dashed arrowhead). This signal is located well below OHCs and it would correspond to DCs’ region (Fig. 6H).

These results show that two cell populations, OHC and SC, are suffering apoptosis at this age in the OC of KO mice.

Then, we analyzed cCAS-3 expression in middle-aged mice. At this age we determined IHCs and SGNs degeneration in KO animals [10]. To identify the labeled cell type in SGN, we used two specific markers, TUBB3 for neurons and Kir4.1 for satellite cells. Using cochlear slices, we found cCAS-3 signal in spiral ganglia corresponding to basal and apical cochlear turns for both genotypes (Fig. 7A). In WT tissue, cCAS-3 was colocalizing with TUBB3 in most of the cases while for KO animals it was either colocalizing with TUBB3 or alone (Fig. 7A). In consequence, we investigated its presence in satellite cells. We observed colocalization of cCAS-3 with Kir4.1 in KO animals while this staining was hardly present in WT (Fig. 7B). These results indicate that satellite cells are also in apoptosis in KO mice.

**Figure 7.**
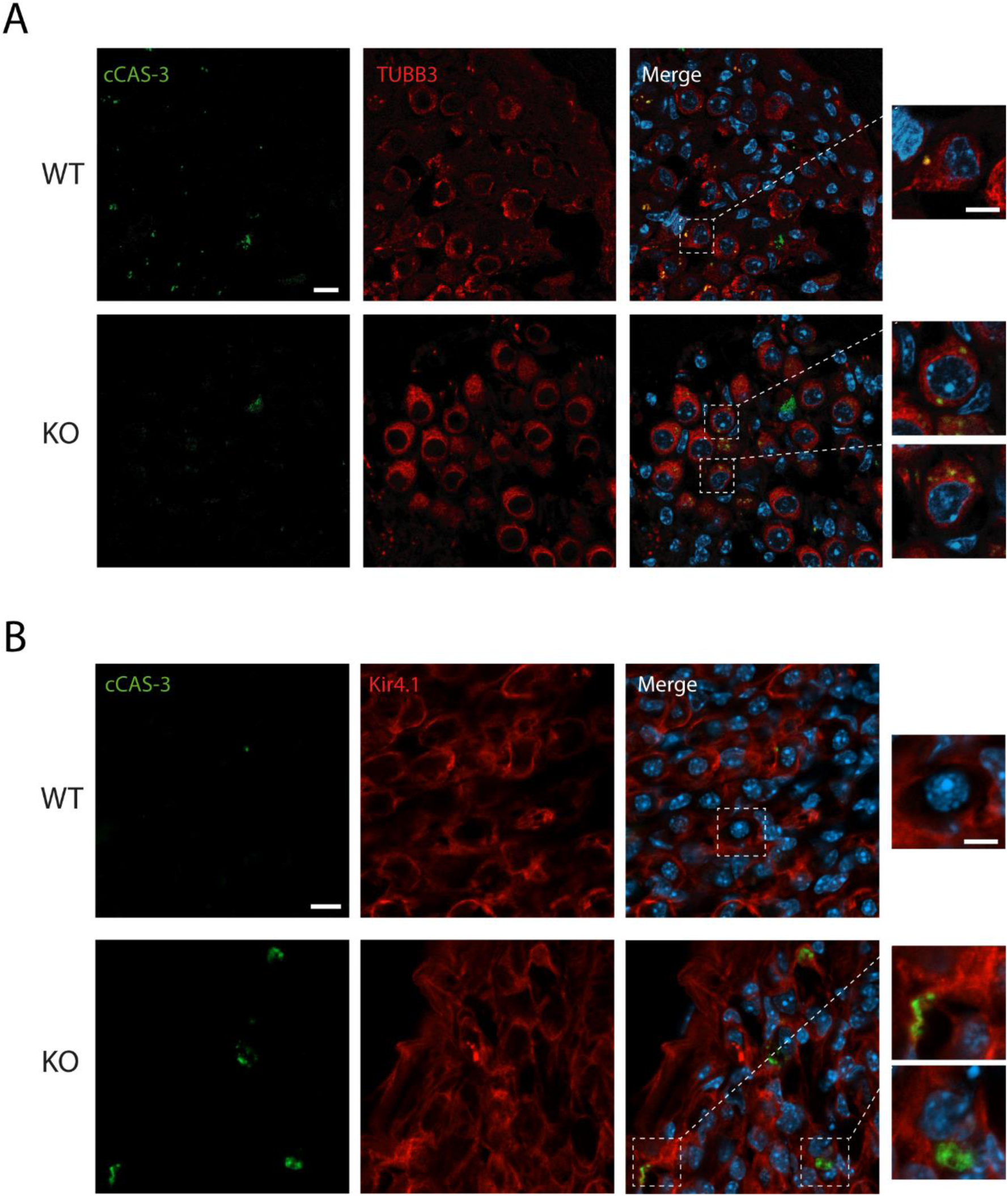
Apoptosis in the spiral ganglia. **A.** Confocal images showing cCAS-3 (green), β-III tubulin (TUBB3, red) and DAPI (blue) staining in middle-aged WT and KO animals. Scale bar: 10 µm (63X). Inset: magnification of the area indicated by dashed-line squares (scale: 2 µm). **B.** Confocal microscopy images showing cCAS-3 (green), Kir4.1 (red) and DAPI (blue) staining in middle-aged WT and KO animals. Scale bar: 5 µm (63X). Inset: magnification of the area indicated by dashed-line squares (scale: 2 µm).

### 3.4. Effects of exposure to loud noise on KCNQ4 KO mouse

As previously described, we found that OHC in KO mice do not function properly due to several tissue alterations. To evaluate if noise exposure induces more damage in the cochlea of this animal model, we performed a sound overexposure to generate an acoustic trauma (AT) test. ABR recordings for each genotype were evaluated at three different time points: control or before trauma (BT), 1 day after AT (1 dAT), and 7 days after AT (7 dAT). ABR measurements showed thresholds increments of 20-40 dB SPL in WT mice 1 dAT (Fig. 8A, compared blue hollow and filled circles), which were statistically significant at frequencies higher than 16.00 kHz, compared to BT (p < 0.0500 for 16.00 kHz, p < 0.0010 for 22.65 kHz, p < 0.0001 for 32.00 kHz and p < 0.0100 for 45.25 kHz; two-way ANOVA, Tukey’s post hoc test). 7 dAT ABR thresholds were still significantly elevated with respect to BT at frequencies higher than 22.65 kHz (p < 0.0500 for 22.65 kHz, p < 0.0010 for 32.00 kHz and p < 0.0100 for 45.25 kHz; two-way ANOVA, Tukey’s post hoc test) (Fig. 8A, light blue circles).

**Figure 8.**
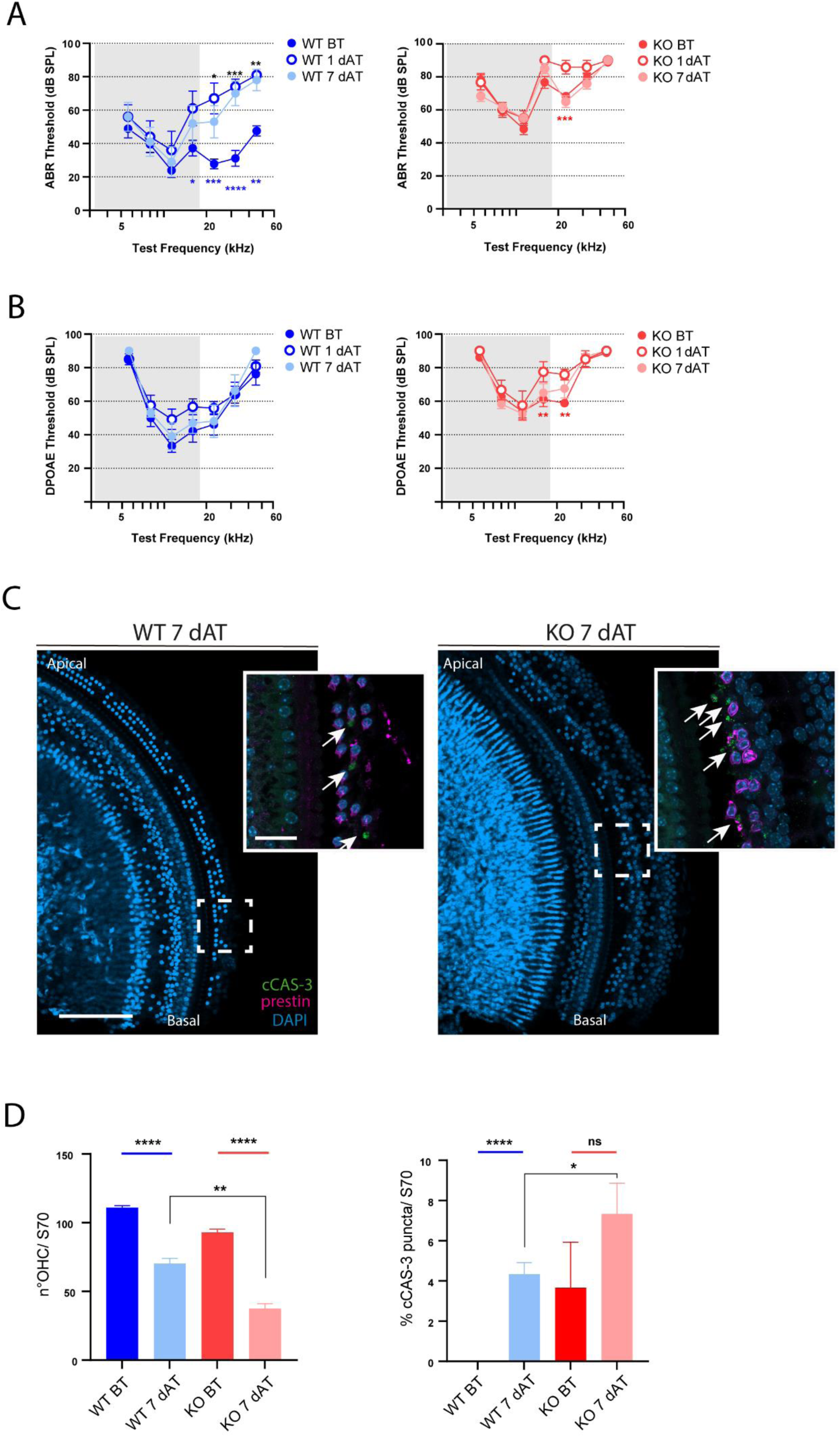
Effects of acoustic trauma on hearing loss in KCNQ4 KO mouse. **A, B. Thresholds after acoustic trauma for ABR (A) and DPOAE (B) from WT and KO mice.** Thresholds were obtained from WT (left) and KO (right) animals beginning at 4 W in a frequency range of 5.60 - 45.25 kHz. Data correspond to animals before trauma (blue and red filled circles, for WT and KO, respectively), 1 day after trauma (1 dAT, blue and red empty circles, for WT and KO, respectively), and 7 day after trauma (7 dAT, light blue and light red filled circles, for WT and KO, respectively). Gray shading represents exposure to 1 - 16 kHz noise at 110 dB for 2 h. Data is presented as mean ± SEM. Statistical analysis was performed using two-way ANOVA, Tukeýs post hoc test. *, p < 0.0500. **, p < 0.0100. ***, p < 0.0010. **C. cCAS-3 staining after acoustic trauma in WT and KO mice**. Images from WT and KO animals of 4 W, showing prestin (red), cCAS-3 (green), and DAPI (blue) staining. White arrows indicate cCAS-3 signal. Scale bar: 100 µm (10X) y 20 µm (inset) (40X). D. Quantification of cCAS-3 signal after acoustic trauma. *Left,* number of OHCs in WT and KO animals before and after 7-day acoustic trauma in the S70 segment of the cochlea (WT BT, blue bar; WT 7 dAT, light blue bar; KO BT, red bar; KO 7 dAT, light red bar). *Right*, cCAS-3 signal quantification in WT and KO animals before and after 7-day acoustic trauma (WT 7 dAT, light blue bar; KO BT, red bar; KO 7 dAT, light red bar). Data are presented as mean ± SEM. Statistical analysis was performed using Student’s t-test. *, p < 0.0500. **, p < 0.0100. ***, p < 0.0010.

In KO animals, auditory thresholds for BT were elevated 20-40 dB compared to those observed for the WT (Fig. 8A). At 1 dAT we determined similar values of auditory thresholds for KO mice compared to BT (Fig. 8A, right, red hollow circles). Only 22.65 kHz exhibited a significant increase in the threshold (p < 0.0010; two-way ANOVA, Tukey’s post hoc test). At 7 dAT all frequencies exhibited similar values as BT (p > 0.0500, two-way ANOVA) (Fig. 8A, right, light red circles). In consequence, AT does not add additional injury to cochlear HCs in the absence of KCNQ4 due to the impairment in hearing transduction. We also determined DPOAE thresholds for WT and KO animals before and after acoustic overexposure (Fig. 8B). For WT we did not find differences between BT and either 1 dAT or 7 dAT. On the contrary, for KO mice we found significant differences between BT and 1 dAT at 16.00 and 22.65 kHz, while at 7 dAT threshold values were similar to BT (Fig. 8B).

After finishing functional studies, mice were euthanized and cochlear tissue was analyzed as previously described. We studied the number of OHCs as well as the presence of the apoptotic marker cCAS-3. Cell survival was analyzed in the S70 region, which is the region with the highest hearing sensitivity. In WT animals, AT generated the loss of OHCs after 7 days of exposure when compared with unexposed mice at the same age (p < 0.0001, Student’s t-test) (Fig. 8C). However, we could not detect loss of IHCs with this protocol (Fig. 8C). OHCs survival decreased around 40 % after 7 dAT (Fig. 8D left, blue vs light blue). In unexposed KO mice we found a 10 % decrease of OHCs compared to WT, which is expected for animals lacking KCNQ4 protein in this region [10]. When we studied KO mice exposed to sound, we found a ∼ 60 % decrease in cell survival, compared to unexposed KO mice (p < 0.0001, Student’s t-test) (Fig. 8D, left, red vs light red). When comparing the percentage decrease at 7 dAT between both genotypes (40 and 60 %, respectively), we did not find significant differences (p > 0.0500, Student’s t-test).

We also analyzed the number of OHCs that expressed cCAS-3 signal. To do this, we counted the number of OHCs in S70 cochlear region and this number was considered the 100 % cell amount. The quantification of cCAS-3 signal was estimated related to this amount. In unexposed animals, cCAS-3 signal was absent in WT animals while, as described previously, it was around 4 % in KO cochleae (Fig. 8D, right). 7 dAT, in WT animals the cCAS-3 signal increased to 4 %, while in KO mice, this amount duplicated in OHCs (Fig. 8D, right). The magnitude of change that was observed for WT and KO animals, related to their unexposed control conditions, was similar, of around 4 %. In consequence, noise trauma did not add a higher damage to OHCs in KO animals.

## 4. DISCUSSION

This study explored the molecular and cellular consequences of KCNQ4 deficiency in the cochlea, providing insights into the progression and mechanisms of HL associated with the expression and function of the KCNQ4 channel. In humans, mutations in this potassium channel lead to HL starting in adolescence, a condition known as DFNA2. In these cases, heterozygous dominant-negative mutations in KCNQ4 drastically impair the extrusion of K^+^ from OHCs. In mouse models of this disease, OHC dysfunction and cell loss have been observed [6, 15, 60], leading to the general understanding that chronic depolarization generates cell dysfunction and eventually triggers cell death [6]. Our findings emphasize the critical role of KCNQ4 in maintaining auditory function and cochlear integrity and reveal new components involved in the development of tissue degeneration.

### 4.1. Functional Studies

Our functional assessments revealed that KCNQ4 deficiency causes progressive auditory dysfunction, evidenced by elevated ABR and DPOAE thresholds across all tested frequencies. KCNQ4 is highly expressed in the basal pole of OHCs, and its absence leads to chronic depolarization, ultimately leading to cell death [6, 7]. However, the progression of this degenerative process begins in the basal turn at around 3 weeks of age [10]. Moreover, the rate of cell loss varies across the cochlear length. Since DPOAE evaluates OHC function [50, 61, 62], our results indicate that nearly all OHCs regardless of their cochlear location, are dysfunctional. However, auditory thresholds, which also reflect OHC amplification function [50, 63], are more elevated in the basal region than in the middle and apical regions. In the basal region, the number of OHCs is already decreased due to cell death by this age [6, 10]. Thus, the severe decline in auditory function reflects not only OHC dysfunction but also cell loss. While young KO mice showed increased auditory thresholds with region-specific sensitivity, middle-aged KO mice exhibited profound deafness, indicating an age-dependent worsening of auditory function as previously reported [10]. This situation would be related to OHC loss in middle and apical turns. The progression of cell death has also been observed in the absence of other potassium channels involved in efferent modulation by the CNS such as BK and SK2 channels, and also in the absence or dysfunction of prestin, the motor protein of OHCs [17, 22, 64]. On the other hand, a similar pattern of cell dysfunction, detectable through functional assays before the loss of OHCs, has been found in the SAMP8 mouse, which suffers HL [65] and in a mutant for mTOR2 [66]. Notably, an important decrease in KCNQ4 expression has been detected in the SAMP8 mouse, highlighting the role of this channel in cell function and survival.

To assess the relationship between KCNQ4 expression and the effects of hearing overstimulation, we analyzed KO and WT animals following acoustic overexposure [67, 68]. The analysis was conducted 7 days after exposure, to minimize contamination of the results due to OHC loss that develops fast in KCNQ4 KO mice at this age [10]. At 1 dAT, we observed a shift in ABR thresholds for frequencies above 20.00 kHz which did not recover to BT values by 7 dAT, suggesting a permanent threshold shift (PTS) [69]. In KO mice, there was no irreversible shift in ABR thresholds at 7 dAT. However, it is important to note that these mice already exhibited an ABR threshold shift prior to acoustic overexposure.

Although PTS is associated with some degree of OHC loss, unexpectedly, DPOAE thresholds in WT animals did not show significant changes after acoustic overexposure despite OHC loss. Nevertheless, studies in different animal models have reported that shifts in DPOAE thresholds do not always correlate with cell loss [70]. While the loss of OHCs typically results in diminished DPOAEs, some studies suggest that residual OHCs can adapt through mechanisms such as prestin upregulation, potentially mitigating the impact of OHC loss on auditory function [71]. This variability is likely influenced by the noise exposure protocol including factors such as exposure duration, distance from loudspeakers or even the genetic background of the mice. For example, C3H/HeJ is a strain reported to be resistant to acoustic trauma [72]. In contrast, IHC functionality in WT mice seems to be more sensitive to acoustic trauma in this strain, possibly due to excitotoxicity, stereocilia damage and loss of synaptic contacts, as reflected by shifts in ABR thresholds previously observed in other strains [73].

### 4.2. Alterations in OHCs

Our functional results prompted us to evaluate structural alterations in this cell type. The proper function of HCs depends on the precise and localized expression of key proteins, such as channels, receptors, and transporters. These cells mobilize K^+^ and Ca^2+^ across their membranes to fulfill their role in the auditory process [5, 17]. OHCs exhibit distinct plasma membrane domains: apical, lateral and basal [17]; which would be generated by lateral wall structures [74]. KCNQ4 is confined in the basal domain after the onset of hearing [7, 19, 24] while prestin localizes exclusively in the lateral domain [75]. Our results showed that in KCNQ4 KO mice, prestin is expressed not only in the lateral domain but also extends into the basal domain. Furthermore, we found mislocalization of the BK channel, which is normally restricted to basal domain [18]. Conversely, similar results on KCNQ4 localization were reported by Takahashi et al., using a mouse model that lacks prestin expression [17]. They observed KCNQ4 mislocalization as a consequence of prestin absence. The overall alterations severely impair OHC function and survival. In WT tissue, just before the onset of hearing, KCNQ4 and prestin mutually exclude each other in the OHC membrane [17]. The mechanism that produces this protein confinement is not totally understood but it would be gated by prestin immobilization on the lateral domain [74]. However, alterations in the expression of any essential protein that conveys the physiological characteristics of OHCs lead to membrane protein disorganization [17, 18, 23, 64]. The evidence presented here would suggest that KCNQ4 is also another key component in establishing membrane compartmentalization and suggest that there could be an additional common factor that links all these protein changes in the disorganization of membrane domains.

In consequence, the modification in key proteins compartmentalization, together with hair bundle alterations that we described previously [10] would impair their amplificatory function that we determined by ABR and DPOAEs measurements.

We also found that the afferent component of the presynapse in OHCs is altered. These cells have ribbon-type synapses that are not as developed as in IHCs [51]. Generally, there is between one to two ribbons in these cells [52, 76–79], and it has been observed that not all ribbons connect with a GluA2 glutamate receptor [78]. In accordance with previous publications, our results confirm the presence of one to two ribbons per OHC. The function of this synapse is not yet clearly understood, and it is thought that they participate in signaling tissue damage induced by loud noises. However, their activation has also been determined in response to sound stimuli of normal intensity that do not generate acoustic damage [80–82]. We found that the number and volume of the ribbon elements were considerably increased. This process has also been observed in mice exposed to acoustic trauma [52]. Although we did not analyze the cellular localization of OHC ribbons and the afferent synaptic boutons, it has been reported that ribbons are located synaptically in the basal region of OHCs and extrasynaptically in the upper medial region of the nucleus equator [77, 79]. In mouse models KO for peripherin or the secreted protein C1LQ1, which exhibit postsynaptic afferent deficiencies, ribbon location and size were not altered [77, 79]. However, another KO mouse model for peripherin showed alterations in ribbon distribution [76]. As these OHC alterations are observed in two different models, this mechanism may represent an adaptation to the increased excitability that OHCs experience during traumatic events.

As we show here, KCNQ4 is necessary not only for K^+^ extrusion but also contribute to proper cell compartmentalization. Furthermore, KCNQ4 is necessary for OHC survival. In KCNQ4 KO mice, OHCs are lost with age due to its absence [6, 10]. It has also been determined that the reduction in KCNQ4 channel expression and function during aging [13, 65, 83, 84] or following acoustic trauma [11, 12, 85] is associated with OHC loss and hearing defects. Proper localization of KCNQ4 in the basal region is necessary to establish the observed electrophysiological properties of the potassium current mediated by KCNQ4, *I_Kn_*, in mature OHCs [86]. In prestin KO mice, KCNQ4 does not localize in the basal region and the absence of clustering was evident in the rightward shift of the activation curves. Additionally, they showed a lower expression of the KCNQ4 channel [86]. These changes in KCNQ4 localization and protein-protein interactions would likely generate a sustained depolarization in OHCs, similarly to what was determined for KCNQ4 KO mice and which may impact cell survival in both models.

Our acoustic overexposure protocol induces the loss of OHCs but not IHCs in WT animals, indicating a PTS [73]. In KO animals, which already show some cell death at 4 W, there was also an increase in the number of absent OHCs. The percentage of cell death compared to the BT condition did not significantly differ between both genotypes. This suggests that acoustic trauma does not cause additional damage to cells that already experience pre-existing tissue stress due to the absence of KCNQ4, which leads to chronic depolarization. As mentioned in the previous section, a loss of OHCs is not always directly related to an increase in DPOAE threshold. Research in rat models has shown that ABR and DPOAEs may appear normal with up to 30% OHC loss [87].

### 4.3. Alterations in IHCs

One of the most significant findings of this study is the structural alterations not only in OHCs but also in IHCs, which could contribute to the hearing loss observed in this animal model. KCNQ4 is also expressed in IHCs. Its presence has been demonstrated at the gene transcription and protein levels, as well as its functional role in stabilizing the membrane potential [88–90]. Its expression is localized in the neck region of the IHC, below the cuticular plate in the apical region. Interestingly, it is also expressed in a region similar to BK [18, 22, 91, 92]. Together with other potassium channels, KCNQ4 contributes to the generation of the three main potassium currents of the IHCs, which provide characteristic properties to the receptor potential of IHCs [88]. Previously, it was found that IHCs from KCNQ4 KO animals exhibit a slight membrane depolarization [6], and we found that these cells degenerate and begins to be lost from 40 W onwards [10]. In the present study, we found structural abnormalities in the IHCs of young animals, leading to IHC dysfunction and contributing to a decline in auditory sensitivity across all frequencies. We observed abnormalities at both the tissue and cellular levels.

We showed that IHC hair bundles exhibited severe alterations in young animals, including stereocilia fusion, elongation, and other abnormalities. Stereocilia fusion may impact auditory signal transduction in IHC by potentially reducing stereocilia movement, as recently reported [93]. Additionally, tip links may be lost, and the number or distribution of MET channels would be affected, ultimately impairing signal transduction. These abnormalities were found along the entire cochlear length. In young KO animals, no loss of IHCs was observed. The presence of stereocilia abnormalities without damage or degeneration of the IHC body suggests the existence of an early “stereociliopathy” [94, 95]. In our previous work, we found early alterations in OHC hair bundle. Alterations restricted to the hair bundles of either IHCs or OHCs have been reported in different rodent models and models of hearing loss associated with aging or noise exposure [93, 94, 96]. In our model, defects in both IHC and OHC hair bundles coexist. At this stage, IHC remain intact, while OHCs degenerate in the basal turn but not in the middle or apical turns [10]. A common functional defect in both cell types is chronic depolarization [6], which is more pronounced in OHCs, as they exhibit more severe stereocilia alterations.

Our functional studies revealed a reduction in auditory sensitivity across all cochlear regions, despite the absence of IHCs loss. These changes partially correlate with the loss of OHC function. However, the magnitude of the changes observed in auditory thresholds and the P1 input-output curves in certain cochlear regions (Fig. 1 and 2) cannot be solely attributed to OHC dysfunction or loss. One of the earliest structural alterations in IHCs under physiological stress, such as loud noise exposure or aging, is synaptic disconnection [97, 98]. As cochlear tissue from KO mice experiences chronic membrane depolarization, we analyzed the integrity of IHC synaptic structures. We found that IHCs displayed significant synaptic disconnection, characterized by a reduction in CtBP2 and GluA2 puncta, consistent with impaired afferent signaling. Similar findings have been reported in other models of hearing loss induced by genetic factors or acoustic trauma [99–101]. Moreover, afferent synaptic loss in IHCs can occur independently of HCs loss or injury, without causing a measurable increase in auditory thresholds (hearing decrement) [98]. Such “synaptopathy” is now considered to be an important early step in age-related pathology [102]. Although early synaptopathy is not the primary feature in our model, our findings suggest that elevated auditory thresholds result from multiple factors beyond OHC dysfunction alone. These structural alterations likely contribute to deficits in auditory processing and signal transmission.

Beyond the reduction in synaptic contacts, we also observed a decrease in ribbon size and number, which could be linked to alterations in the actin filament network that is in close contact with ribbons. We detected a reduction in MYO7A intensity in IHCs (data not shown). MYO7A is highly localized in the cuticular plate and stereocilia but is also found in the cytoplasm, mainly in the subnuclear region of IHCs [103]. Actin filaments network are known to regulate ribbon synapse function, as previously reported [104, 105]. Recently, Cortada et al. [66] described structural alterations in both the hair bundle and the actin cytoskeleton surrounding IHC synapses in their model, proposing that these changes contribute to synapse loss.

Despite Kharkovets et al. reported that the afferent synaptic zone appears normal in KO animals of mixed genetic background at 4 months of age [6], we found alterations in the number and volume of afferent synapses in IHC, in agreement with our functional studies mentioned above. These differences could be generated by the different genetic background that we employed for our experiments.

Using our acoustic overexposure protocol, we did not observe a significant IHC loss. This has also been reported by other authors, who determined that prolonged exposure to loud noise is necessary to observe IHC and SC death [106]. In our KO mice, we also did not observe IHC loss due to acoustic overexposure. At this age, there is no IHC loss associated with the absence of KCNQ4 either [10].

### 4.4 Mechanisms of Cell death

Here, we reveal that apoptosis is a key mechanism underlying the loss of OHCs in KO animals. Activation of this pathway was observed as early as 4 W in the basal cochlear turn, consistent with the onset of cellular degeneration observed in cytocochleograms analyses [10]. As aging progresses, the presence of the cCAS-3 signal expands to the middle and apical turns, eventually affecting the entire cochlear length. Moreover, gene expression analysis revealed elevated *Bax* mRNA levels and reduced *Bcl-2* expression, resulting in an increased *Bax/Bcl-2* ratio, which indicates enhanced intrinsic apoptosis pathway activation [34, 99]. Apoptosis has been extensively studied in various hearing impairment processes. Mutations in apoptosis-associated genes, such as DFNA5 and DFNB74, have been linked to hearing loss [34]. Additionally, ototoxic drugs, including aminoglycoside antibiotics and cisplatin, are known to trigger apoptotic pathway [107, 108]. In the context of NIHL, apoptosis is a well-established contributor to cochlear damage. Multiple animal models studies have explored different pathways through which apoptosis is triggered [109, 110]. Our results align with these findings, indicating that in an environment of chronic depolarization, apoptosis is one of the pathways involved in cochlear cell death.

We also determined that apoptosis occurs in DCs from young KO mice. Although there are limited references documenting SCs loss in HL models, our data highlight their vulnerability alongside OHCs [37]. Apoptosis in cochlear SCs, particularly in Kölliker’s organ, plays a significant role in both cochlear development and pathology [38, 111]. However, we could not determine whether this process in SCs occurs before or after OHC death. In this context, another study observed that DCs apoptosis leads to OHC loss [112].

Apoptosis also contributes to the loss of SGNs in KO animals. The cCAS-3 signal was detected both in the neuronal cytoplasm and surrounding the SGN soma, which is encased by glial cells known as satellite cells [113]. These cells, along with Schwann cells, support neuron metabolism and regeneration [114]. In the KO cochlea, the apoptotic signal around SGNs likely indicates apoptosis of satellite cells, which were identified with Kir4.1, a specific marker.

Concerning neurons, lack of trophic support, possibly from IHCs or surrounding glial cells, is suggested as a cause of SGN death [115]. When IHCs degenerate due to noise exposure or ototoxic drugs, SGN death follows as a result of the loss of trophic support [116]. The majority of IHC afferent innervation comes from SGNs, making this link plausible [117]. Additionally, synaptic alterations in IHCs have been observed, further supporting the hypothesis of synaptic changes contributing to SGN death. [118–120].

## 5. CONCLUSION

This study enlightens the essential role of KCNQ4 in preserving cochlear function and auditory health. It reveals that cochlear tissue alteration begins early in life. The absence of KCNQ4 disrupts the localization of key proteins, such as prestin and BK channels, and also leads to structural abnormalities, including stereocilia defects and synaptopathy, further compromising HCs functionality. Together, all these changes observed in the cells that make up the cochlear tissue induce chronic cellular stress, which, either directly or indirectly, leads to apoptosis and a worsening of auditory function with age. These findings highlight the significance of KCNQ4 in maintaining cochlear function and suggest potential pathways for therapeutic intervention. Future research targeting these mechanisms may enhance strategies to prevent or mitigate hearing loss associated with KCNQ4 dysfunction.

## Acknowledgements

ER, CC, VC, GP have a doctoral fellowship from the Argentina National Research Council (CONICET-UNS). IO had an advance student fellowship from the National South University.

## CRediT authorship contribution statement

Ezequiel Rías: Conceptualization, Formal Analysis, Investigation, Methodology, Software, Resources, Validation, Visualization, Writing-original draft, Writing-review and editing.

Camila Carignano: Conceptualization, Formal Analysis, Investigation, Methodology, Software, Resources, Validation, Visualization, Writing-original draft, Writing-review and editing.

Valeria C. Castagna: Formal Analysis, Investigation, Methodology, Software, Resources, Validation.

Leonardo Dionisio: Conceptualization, Formal Analysis, Funding Acquisition, Investigation, Methodology, Project Administration, Resources, Supervision, Validation, Visualization, Writing-original draft, Writing-review and editing.

Jimena A. Ballestero: Formal Analysis, Investigation, Methodology, Software, Resources, Validation.

Giuliana Paolillo: Investigation, Methodology, Resources.

Ingrid Ouwerkerk: Formal Analysis, Investigation, Methodology, Software, Resources, Validation.

María Eugenia Gómez-Casati: Conceptualization, Formal Analysis, Investigation, Resources, Supervision, Validation, Visualization, Writing-original draft, Writing-review and editing.

Guillermo Spitzmaul: Conceptualization, Formal Analysis, Funding Acquisition, Investigation, Project Administration, Resources, Supervision, Validation, Visualization, Writing-original draft, Writing-review and editing.

---

**Supplementary Figure 1.**
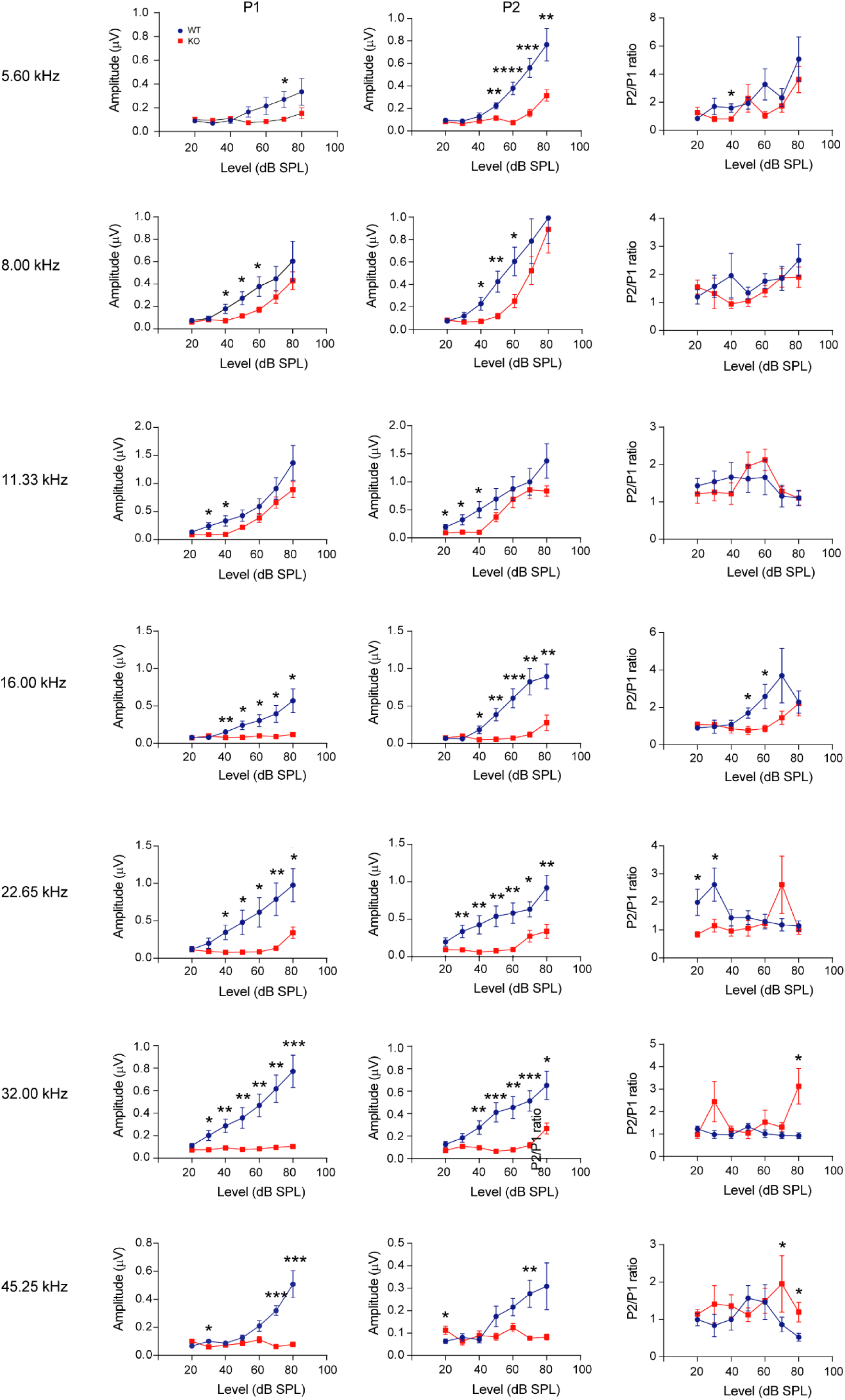
Peak 1 (P1), Peak 2 (P2) and P2/P1 ratio analysis. P1 and P2 values from WT and KO animals of 4 W in a frequency range of 5.60 - 45.25 kHz and in an intensity range of 20 - 80 dB SPL. Data are presented as mean ± SEM. Statistical analysis was performed using Student’s t-test between WT and KO at each intensity (n = 9). *, p < 0.0500. **, p < 0.0100. ***, p < 0.0010, ****, p < 0.0001.

**Supplementary Figure 2.**
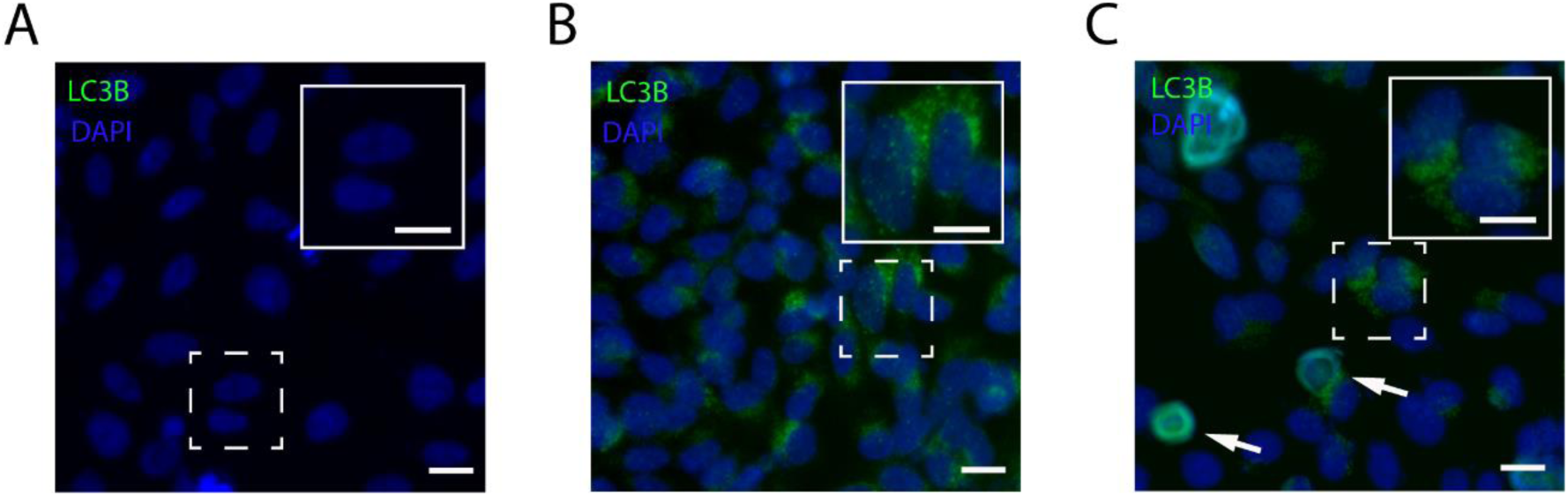
Autophagy testing in the D407 cell line using LC3B staining. Two different treatments were used to test the autophagy pathway: deprivation of fetal bovine serum (FBS) and incubation with 150 µM of H_2_O_2_ for 17 h. **A**. Microscopy image of LC3B staining (green) and DAPI (blue) of D407 cell culture without any treatment (control). **B**. Microscopy image of LC3B staining (green) and DAPI (blue) of D407 cell culture during deprivation of FBS. **C.** Microscopy image of LC3B staining (green) and DAPI (blue) of D407 cell culture during deprivation of FBS and H_2_O_2_ incubation. White arrows indicate cells with rounded shape and positive LC3B staining (**C**). White dashed boxes indicate the area of magnification. Scale: 10 µm.

